# A trans-amplifying mRNA vaccine with consensus spike elicits broadly cross-neutralizing antibody response against multiple SARS-CoV-2 variants

**DOI:** 10.1101/2025.01.22.634352

**Authors:** Abhinay Gontu, Sougat Misra, Shubhada K. Chothe, Santhamani Ramasamy, Padmaja Jakka, Maurice Byukusenge, Lindsey C. LaBella, Meera Surendran Nair, Bhushan M Jayarao, Marco Archetti, Ruth H. Nissly, Suresh V. Kuchipudi

## Abstract

SARS-CoV-2 continues to evolve and evade vaccine immunity necessitating vaccines that offer broad protection across variants. Conventional mRNA vaccines face cost and scalability challenges, prompting the exploration of alternative platforms like trans-amplifying (TA) mRNA that offer advantages in safety, manufacturability, and antigen dose optimization. Using consensus sequences of immunodominant antigens is a promising antigen design strategy for board cross-protection. Combining these two features, we designed and evaluated a TA mRNA vaccine encoding a consensus spike protein from SARS-CoV-2. Mice receiving the TA mRNA vaccine produced neutralizing antibody levels comparable to a conventional mRNA vaccine using 40 times less antigen mRNA. In hACE2 transgenic mice challenged with the Omicron BA.1 variant, the TA mRNA vaccine reduced lung viral titers by over 10-fold and induced broadly cross-neutralizing antibodies against multiple variants. These findings highlight the potential of TA mRNA vaccines with consensus antigen design, to improve efficacy and adaptability against SARS-CoV-2 variants.

## Introduction

Since its emergence in 2019, Severe Acute Respiratory Syndrome Coronavirus 2 (SARS-CoV-2) has undergone continual genetic evolution, leading to the emergence of multiple variants^1,2^. Certain variants of SARS-CoV-2 are designated as Variants of Interest (VOI), Variants of Concern (VOC), Variants of High Consequence (VOHC), or Variants Being Monitored (VBM) based on specific attributes and characteristics that might necessitate public health interventions^3,4^. VOCs have displayed increased transmissibility, and virulence, reduced susceptibility to neutralization by antibodies from natural infection or vaccination, evasion of detection, or compromised efficacy against therapeutics^5, 6^. As of January 2025, five VOCs have been recognized: Alpha, Beta, Gamma, Delta, and Omicron, with notable differences in transmissibility and capacity to escape vaccine immunity^3^. While multiple factors contribute to the evolution of SARS-CoV-2^7, 8^, some of the key contributors include the infidelity of the RNA polymerase, recombination, host immune pressure, and the virus circulation in animal reservoirs^1, 9^.

Vaccination remains the most effective strategy to prevent severe infection, hospitalization and death from Coronavirus disease-2019 (COVID-19), caused by SARS-CoV-2^10, 11^. However, the emergence of novel variants that compromise vaccine immunity presents a significant challenge, requiring continuous evaluation of vaccine efficacy, and regular updates to maintain effectiveness^12^. The first-generation COVID-19 vaccines were designed using the spike (S) protein from the ancestral SARS-CoV-2 B.1 lineage^13^. Vaccine formulations based solely on the ancestral variant offered reduced protection against VOCs, particularly Beta, Gamma, Delta, and Omicron^14, 15, 16^. mRNA vaccines have rapidly undergone updates to confer protection against circulating variants of SARS-CoV-2, showcasing their effectiveness in addressing pandemics^17^. While updated booster shots have been developed to include additional S protein mRNAs to match circulating variants, they are often less effective since new variants have already emerged by the time, they are ready for deployment. Additionally, the limited coverage of booster shots post-primary vaccination contributes to vaccine ineffectiveness, leaving a significant portion of the global population vulnerable to reinfection^18^.Hence, there is a critical need to design antigens that provide broad-spectrum protection across SARS-CoV-2 variants. The existing mRNA vaccine formulations also face challenges of inadequate global supply, primarily stemming from the need for a high mRNA load to encode the spike protein of SARS-CoV-2^19, 20, 21^. Therefore, it is crucial to develop next-generation mRNA vaccines, that offer broad protection against various SARS-CoV-2 variants while requiring lower doses reduced dosage requirements.

Amplifying mRNA vaccines, distinguished by their dose-sparing attributes, offer an attractive solution to overcome limited supplies of conventional mRNA vaccines^22, 23^. These vaccines carry their replicase (virus-encoded RNA-dependent RNA polymerase), facilitating amplification of the gene of interest following delivery into cells^24, 25^. Two primary categories of amplifying mRNA vaccines are self-amplifying (SA mRNA) and trans-amplifying (TA mRNA)^24, 25^. The key distinction lies in the administration of replicase, with TA mRNA vaccines delivering it as a separate mRNA (trans) from the one coding for the gene of interest, while SA mRNA vaccines deliver the replicase on the same mRNA (cis or self) as the gene of interest. Few studies have demonstrated the dose-sparing characteristics of SA mRNA vaccines for SARS-CoV-2 in animal models and human clinical trials^26, 27, 28^. Nevertheless, SA mRNA vaccines have limitations, such as the limited flexibility to accommodate nucleoside modifications, the substantial length of the construct affecting manufacturability, and the inability to be multiplexed^25^. In contrast, the TA mRNA approach offers greater modification flexibility, allowing for multiplexing and enhancing the intrinsic versatility and scalability of mRNA vaccine technologies^23, 29^. Additionally, the TA mRNA approach has demonstrated increased translation efficiency of mRNA vaccines, suggesting it may solve the current challenges posed by SARS-CoV-2^23^.

The S protein is the primary antigen in all major SARS-CoV-2 vaccines^30^. Existing SARS-CoV-2 vaccine formulations have employed the S protein sequence derived from clinical isolates, with minimal modifications aimed at enhancing protein expression and stability^31^. To improve vaccine efficacy, the adoption of a consensus spike protein sequence, synthesized using amino acid sequences conserved across multiple variants of SARS-CoV-2, represents a promising strategy for achieving comprehensive immunity. Vaccines based on consensus sequence antigen design have demonstrated success in addressing various viral infections, including Influenza, HIV, and Dengue^32, 33, 34^. Additionally, vaccine formulations based on consensus antigen design have shown broad cross-protection against other coronaviruses like Avian coronavirus and MERS-CoV^35, 36^, making this design a promising strategy for next-generation SARS-CoV-2 vaccines.

To address the challenges of current vaccine strategies for SARS-CoV-2, we designed and evaluated a TA mRNA vaccine based on a consensus spike protein sequence, with the goal of developing a vaccine capable of generating robust, neutralizing immune responses across a broad range of SARS-CoV-2 variants while reducing the required antigenic dose. The immunogenicity of the TA mRNA vaccine formulations were rigorously evaluated in hACE2-expressing transgenic mice, offering promising, dose-sparing alternatives to conventional mRNA vaccines. Additionally, this study investigated the replication dynamics of mRNA within the TA mRNA system using an epithelial cell model to gain deeper insights into its mechanism of action.

This approach holds the potential to pave the way for next-generation vaccines that offer more robust, efficient and broadly cross-protective immunity against emerging SARS-CoV-2 variants.

## Results

### Design of consensus SARS-CoV-2 spike antigen

A consensus sequence for the SARS-CoV-2 spike antigen was designed by aligning nine SARS-CoV-2 variants using Multiple Sequence Comparison by Log-Expectation (MUSCLE). The analysis revealed a high degree of sequence similarity among earlier variants (ancestral B.1 lineage to Delta), with percentage identities of 98.82% to 98.35%. The newer Omicron variants (BA.1 through BA.5) displayed greater amino acid variation, resulting in a lower percentage similarity of 96.63% to 97.17%. **Fig. 1A** provides a detailed comparison of the spike protein sequences, including both the consensus sequence and the nine variants, highlighting the similarities and differences in amino acid sequences.

**Fig 1.**
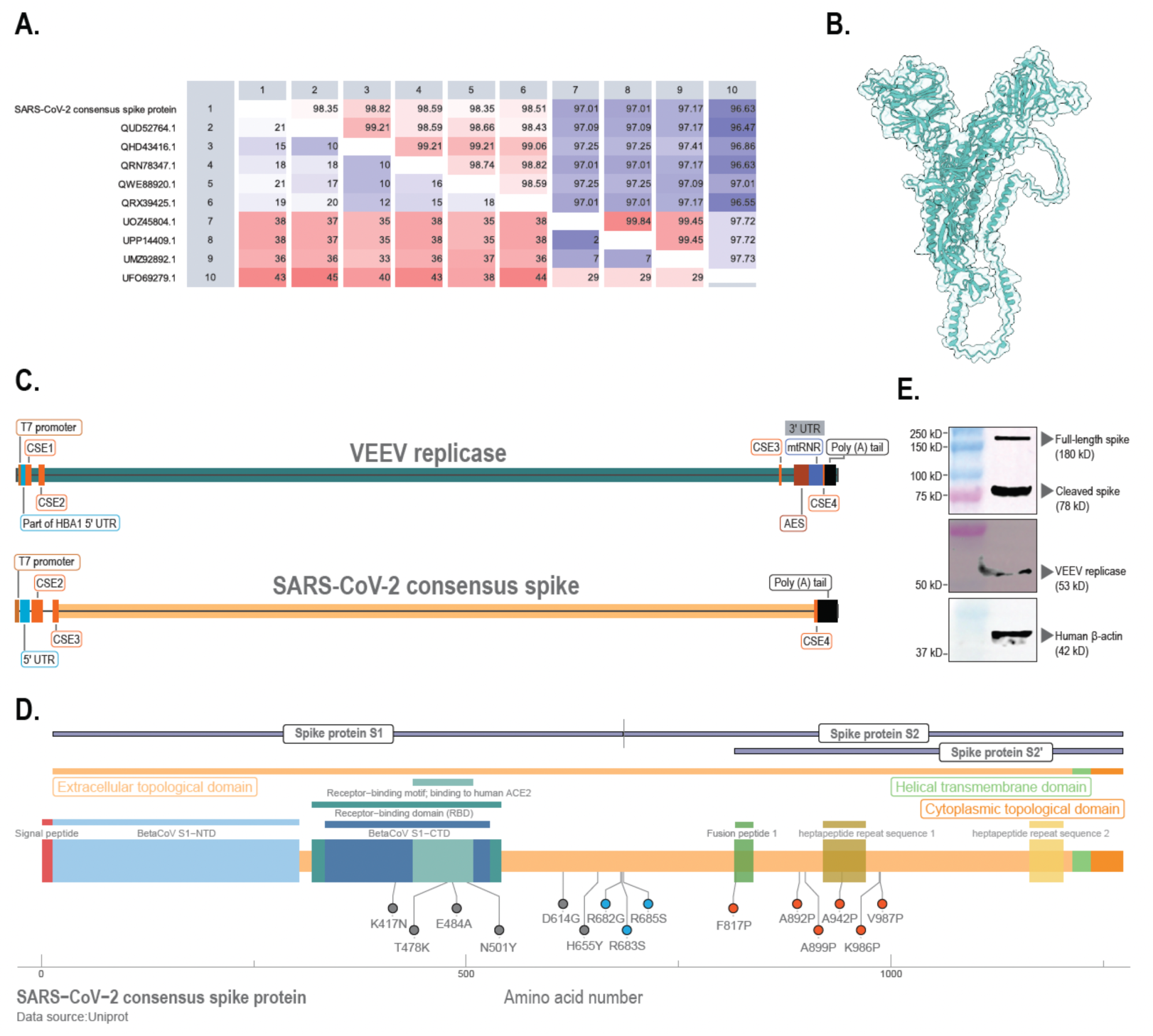
Design aspects and protein expression of mRNA constructs in the trans-amplifying mRNA (TA mRNA) vaccine coding for SARS-CoV-2 consensus spike. **(A)** The table shows pairwise comparisons between SARS-CoV-2 spike protein sequences, comprising a consensus sequence and nine variants. The upper-right triangle (blue) displays percentage identity values, while the lower-left triangle (white) demonstrates the number of amino acid differences between sequences. The consensus spike used in this study exhibited close similarity (98.82% to 98.35%) to the earlier SARS-CoV-2 variants (ancestral to Delta) and lower similarity (96.63% to 97.17%) to the later variants (Omicron BA.1/BA.2 to Omicron BA.5, as tested in the study). **(B)** Structural superimposition of the SARS-CoV-2 consensus spike protein (white) with the SARS-CoV-2 spike protein 6XKL (teal) was performed. The sequence alignment score was 5565.7, and the Root Mean Square Deviation (RMSD) between 745 pruned atom pairs was 1.154 Å, indicating high structural similarity. **(C)** The TA mRNA vaccine comprises two mRNA constructs: one encoding the replicase derived from the Venezuelan Equine Encephalitis Virus (VEEV), preceded by the 5’UTR from the HBA1 gene and followed by the 3’UTR from AES and mtRNR1; and another encoding the SARS-CoV-2 consensus spike protein, with sequences flanking it coding for conserved sequence elements (CSEs) of the VEEV replicase. **(D)** A schematic representation of the SARS-CoV-2 consensus spike protein highlights the sequence modifications relative to the ancestral B.1 lineage spike protein (GenBank: QHD43416.1). Modifications shown in grey represent consensus changes derived from variant alignments. Additional modifications were introduced to stabilize the spike protein in its pre-fusion conformation: three alterations at the S1/S2 cleavage site (shown in green) and six proline substitutions (shown in red). **(E)** A549 cells were transfected with mRNAs encoding the TA mRNA vaccine. Four hours post-transfection, total protein was subjected to western blot analysis using specific antibodies to detect protein expression. The consensus spike protein was observed in both full-length (180 kDa) and cleaved (78 kDa) forms, while the replicase appeared as a 53 kDa protein. Beta-actin (42 kDa) served as a reference control for the cells (Lane 1: Marker, Lane 2: Cell lysate)

Structural predictions for the SARS-CoV-2 consensus spike protein were generated using the AlphaFold program. This predicted structure was then compared to the spike protein represented by Protein data bank (PDB) ID 6XKL. The sequence alignment achieved a score of 5565.7, and the Root Mean Square Deviation (RMSD) between the consensus spike protein and the 6XKL structure was 1.154 Å, based on 745 pruned atom pairs. The low RMSD value indicates a high structural similarity between the consensus spike protein and the 6XKL structure, supporting the functional relevance of the consensus spike as a vaccine candidate. **Fig. 1B** illustrates the structural superimposition of the SARS-CoV-2 consensus spike protein with the 6XKL spike protein.

Comparative analysis of the consensus spike protein sequence revealed six amino acid differences relative to the B.1 lineage spike protein. Most of these differences were located in the receptor-binding domain (RBD) and included mutations such as K417N, T478K, E484A, and N501Y. Additional mutations at the S1/S2 cleavage site (D614G) and the furin cleavage site (H655Y) were also noted. To enhance the stability of the spike protein in its pre-fusion state, additional substitutions were introduced at the S1/S2 cleavage site (R682G, R683S, R685S), fusion peptide 1 (F817P), heptapeptide repeat sequence 1 region (A892P, A899P, A942P) and the conventionally used 2P mutations (K986P and V987P) ^37^. **Fig. 1D** depicts all the amino acid changes and substitutions introduced in the consensus spike protein.

### Design of trans-amplifying mRNA vaccine expressing consensus SARS-CoV-2 spike antigen

The TA mRNA vaccine was engineered utilizing two distinct mRNA constructs: one encoding the VEEV replicase and the other encoding the gene of interest. The VEEV replicase elements were strategically flanked at both termini of the mRNA coding for the gene of interest to enhance targeted amplification and replication by the VEEV replicase. In alignment with the methodology proposed by Beissert *et al.* for the validation of the TA mRNA design, we selected nano-luciferase as a quantifiable gene of interest ^23^. Following the transfection of 293T cells with mRNAs coding for nano-luciferase and VEEV replicase, we noted a significant increase (p= 0.0033, t-test) in the luciferase output in cells co-transfected with both the replicase and nano-luciferase compared to those receiving nano-luciferase alone **(Supplementary data.2)**.

Upon successfully validating the design, we formulated SARS-CoV-2 TA mRNA vaccine by incorporating mRNAs coding for VEEV replicase and the consensus spike protein. **Fig. 1C** shows the design of the mRNA constructs utilized in the vaccine. *In vitro* transcription, capping, and purification of the mRNA constructs were conducted, after which the mRNAs were transfected into A549 cells. Cell lysates were collected four hours post-transfection and subjected to analysis via western blot. The consensus spike protein was identified in both its full-length (180 kDa) and cleaved (78 kDa) forms. The VEEV replicase protein was detected as a band at 53 kDa. β-actin (42 kDa) served as the reference control for protein expression analysis. **Fig. 1E** presents the western blot results of the mRNAs used in the vaccine. Subsequently, following the studies by Besisert *et al.* and Schmidt *et al.,* where the TA mRNA vaccine was evaluated *in vitro* at two-time points, we performed western blots at 4 hours and 12 hours following the co-transfection of the TA mRNA constructs (VEEV replicase and Consensus SARS-CoV-2 spike) ^23, 29^. Despite facing limitations with the amount of transfectable input mRNA of SARS-CoV-2 in an *in vitro* cell-based experiment, we demonstrated an increase in the SARS-CoV-2 protein content at the 8-hour time-point **(Supplementary data.4)**.

### mRNA vaccination induces SARS-CoV-2 neutralizing antibody responses in mice

Three distinct vaccine formulations were developed. One vaccine formulation consisted of 20 μg/dose of standard mRNA encoding the SARS-CoV-2 consensus spike protein, while the other two vaccine groups employed trans-amplifying designs. One of the trans-amplifying vaccines contained 0.5 μg/dose of SARS-CoV-2 consensus spike RNA and 20 μg/dose of VEEV replicase mRNA (TA mRNA-high dose or TAHD). The spike mRNA composition in the TA mRNA-high dose is 40 times less than the conventional mRNA vaccine. Another trans-amplifying vaccine formulation (TA mRNA-low dose or TALD) was made with 400 times lesser spike mRNA composition, resulting in 0.05 μg/dose of SARS-CoV-2 consensus spike RNA and 20 μg/dose of VEEV replicase mRNA. The vaccines were tested for immunogenicity in mice. Mice were organized into four groups, each consisting of six animals. Phosphate-buffered saline (PBS) was used for mock vaccination in the control group of mice. All mice were vaccinated twice, with an interval of two weeks between the doses. Antibody analyses were conducted on blood samples collected two weeks after the second dose.

All mRNA-vaccinated mice (Consensus spike mRNA group and TA mRNA groups) developed strong neutralizing antibody responses after two immunizations, with the mock-vaccinated (PBS) mice showing no detectable responses (**Fig. 2, 3**). The log_2_ median live virus neutralization (VN) titers were 7.32 (95% CI: 6.70 to 8.28) for the 20 μg consensus spike mRNA group, 6.82 (95% CI: 6.25 to 7.40) for the high-dose TA mRNA group, and 4.32 (95% CI: 4.11 to 5.20) for the low-dose TA mRNA group (**Fig. 2C**). Statistical analysis with the Kruskal-Wallis test revealed significant differences in serum neutralization titers among the groups (p = 0.0014). The neutralization titers of the high-dose TA mRNA group were comparable to those of the 20 μg consensus spike mRNA group with no significant difference (p = 0.14), while both the consensus spike mRNA and high-dose TA mRNA groups had significantly higher (p = 0.0037 and p = 0.0039, respectively) titers than the low-dose TA mRNA group.

**Fig 2.**
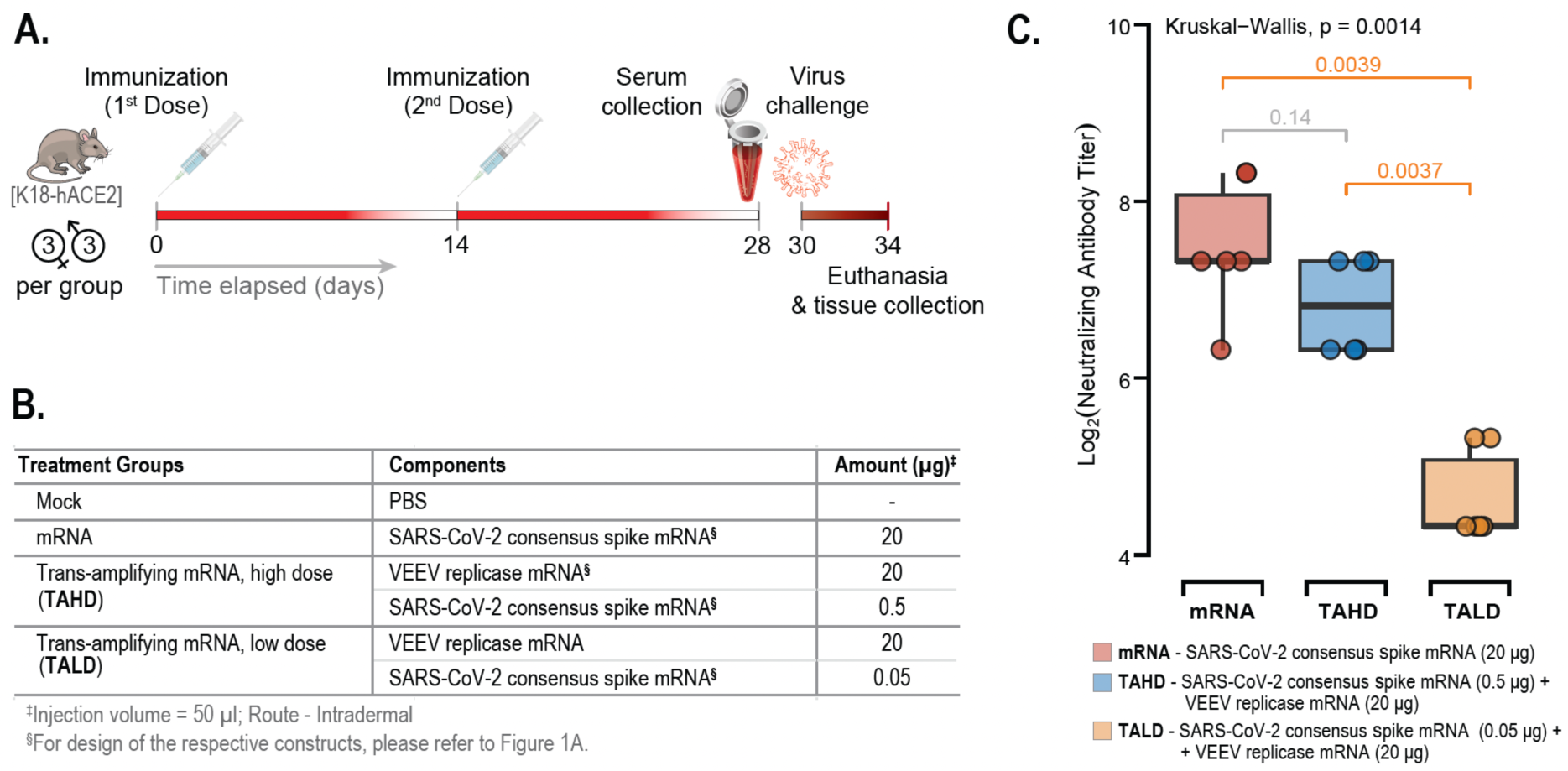
Experimental design and virus neutralization titers in hACE2-expressing transgenic mice immunized with conventional or trans-amplifying mRNA vaccines. **(A)** Overview of the timeline for immunization, blood collection, and challenge procedures in the experimental groups (*n*=6 per group, 3 males and 3 females). **(B)** Description of the treatment groups, including details of the administered vaccines (components, mRNA dosage, volume, and route). **(C)** Results of the virus neutralization assay using the SARS-CoV-2 Delta variant (B.1.617.2) indicated differences between the treatment groups (p=0.0014, Kruskal-Wallis Test followed by Wilcoxon Rank Test for pair-wise comparison). Serum neutralization titers from mice receiving 0.5 μg of consensus spike mRNA (mRNA) with 20 μg of replicase mRNA (Trans-amplifying mRNA: high dose, TAHD) were comparable to those from mice receiving 20 μg of consensus spike mRNA (mRNA) (p=0.14, Kruskal-Wallis Test). Both the TAHD and mRNA groups showed significantly higher titers compared to the Trans-amplifying mRNA: low dose group (TALD) (p=0.0037 and p=0.0039, respectively, Kruskal-Wallis Test). The plot shows individual neutralization titers with median values and 95% confidence intervals. These results demonstrated that TAHD, with an mRNA antigen load 40 times lower, could generate neutralizing antibody titers equivalent to those produced by conventional mRNA without trans-replicase. (Note: Mock-vaccinated mice did not produce any detectable neutralizing antibody responses; data not shown).

**Fig 3.**
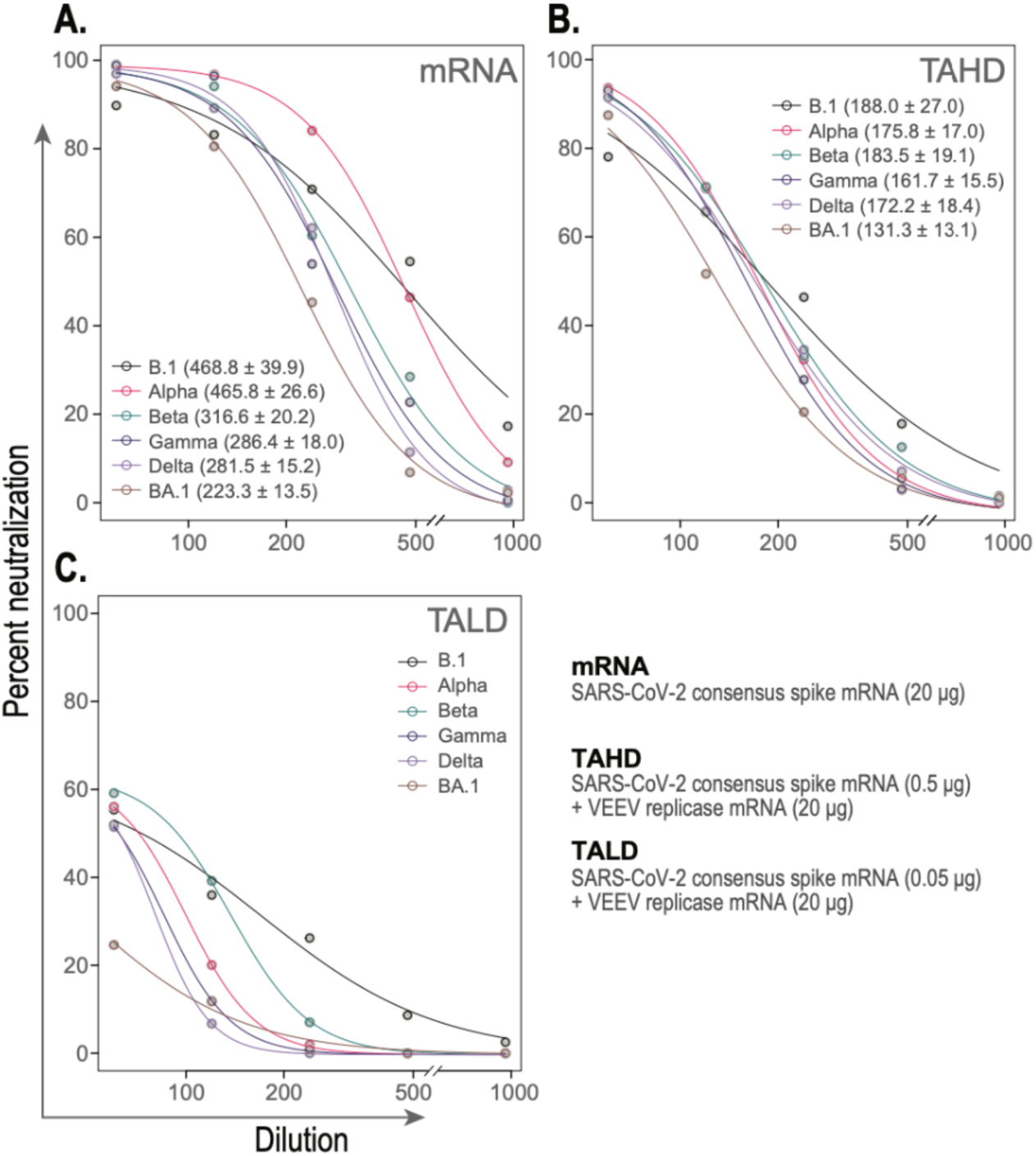
Pseudovirus neutralizing titers of serum samples collected from vaccinated mice. Transgenic hACE2-expressing mice, (*n*=6 per group, 3 males and 3 females), were immunized with different mRNA formulations administered intradermally at two-week intervals. The mRNA group received 20 μg of consensus spike mRNA (mRNA), the Trans-amplifying mRNA: high dose (TAHD) received 0.5 μg of consensus spike mRNA combined with 20 μg of replicase mRNA, and the Trans-amplifying mRNA: low dose (TALD) group received 0.05 μg of consensus spike mRNA with 20 μg of replicase mRNA. Mock-vaccinated mice received an equivalent volume (50 μL of phosphate-buffered saline, PBS). Serum samples were collected on day 28 and subjected to a neutralization assay using various SARS-CoV-2 pseudoviruses. The half-maximal effective concentration (EC50) values were derived from the dose-response curves using a four-parameter Hill equation to assess pseudovirus neutralization efficacy. **(A)** Serum from mice immunized with 20 μg of consensus spike mRNA exhibited antibody titers against all variants, with EC50 values (serum dilution factor) ranging from 223.3 ± 13.5 (Omicron BA.1) to 468.8 ± 39.9 (B.1 ancestral variant). **(B)** Mice immunized with a ‘40-fold lower dose of consensus spike mRNA’ (0.5 μg) in combination with VEEV replicase (TAHD) also demonstrated antibody titers against all SARS-CoV-2 pseudovirus variants, with EC50 values ranging from 131.3 ± 13.1 (Omicron BA.1) to 188.0 ± 27.0 (B.1 ancestral variant). **(C)** Immunization with a low dose of consensus spike mRNA (0.05 μg) combined with VEEV replicase (TALD) resulted in minimal neutralization efficacy. EC50 values were not calculated for this group, as the highest neutralization responses against all pseudovirus variants were not substantial enough to compute EC50 values with a high degree of confidence.

Serum samples with measurable levels of VN titers were assessed for the breadth of their antibody responses through neutralization tests with SARS-CoV-2 pseudoviruses expressing spike proteins from multiple variants. The half-maximal effective concentration (EC_50_) values indicated high neutralization efficacy for the consensus spike mRNA group: 468.8 ± 39.9 (B.1), 465.8 ± 26.6 (Alpha), 316.6 ± 20.2 (Beta), 286.4 ± 18.0 (Gamma), 281.5 ± 15.2 (Delta), and 223.3 ± 13.5 (Omicron BA.1). The high-dose TA mRNA group (0.5 μg spike mRNA) showed more uniform EC_50_ values across variants: 188.0 ± 27.0 (B.1), 175.8 ± 17.0 (Alpha), 183.5 ± 19.1 (Beta), 161.7 ± 15.5 (Gamma), 172.2 ± 18.4 (Delta), and 131.3 ± 13.1 (Omicron BA.1). While neutralization was observed in the low-dose TA mRNA group (0.05 μg spike mRNA), the values were low and the dose-response curves could not be reliably fitted. **Fig. 3** displays the dose-response curves and EC_50_ values for vaccinated mice against SARS-CoV-2 pseudoviruses.

### The trans-amplifying mRNA vaccine achieves comparable efficacy to conventional mRNA vaccines using a substantially lower spike antigen dose

The efficacy of the vaccine formulations in reducing viral load was evaluated by measuring N1 gene copy numbers and infectious virus titers in the lungs of vaccinated and control mice post SARS-CoV-2-challenge. We estimated the SARS-CoV-2 viral load in mice by measuring the N1 gene copy numbers and 50% tissue culture infective dose (TCID_50_) from lung tissue. The Log_10_ median N1 gene copy numbers were 5.50 (95% CI: 5.07 to 6.23) in the mock-vaccinated group, 3.10 (95% CI: 1.85 to 4.30) in mice vaccinated with 20 μg of consensus spike mRNA (2 mice did not show detectable N1 copy number), 4.53 (95% CI: 4.39 to 4.76) in mice vaccinated with TA mRNA-high dose, and 5.11 (95% CI: 4.95 to 5.25) in mice receiving the TA mRNA-low dose (**Fig. 4A**). The Kruskal-Wallis test indicated a significant difference in N1 gene copy numbers among the groups (p = 0.0004). Post-hoc Wilcoxon rank-sum tests revealed that N1 copy numbers were significantly lower in mice vaccinated with consensus spike mRNA (p = 0.0095) and TA mRNA-high dose (p = 0.0022) compared to the mock-vaccinated group. The N1 copy numbers in the TA mRNA-low dose group did not differ significantly from those in the high-dose TA mRNA group (p = 0.065). The N1 copy numbers in the consensus spike mRNA group were significantly lower than those in both the TA mRNA-high dose (p = 0.0095) and the TA mRNA-low dose (p = 0.0095) groups. Additionally, the TA mRNA-high dose group had significantly lower N1 copy numbers compared to the TA mRNA-low dose group (p = 0.0022).

**Fig 4.**
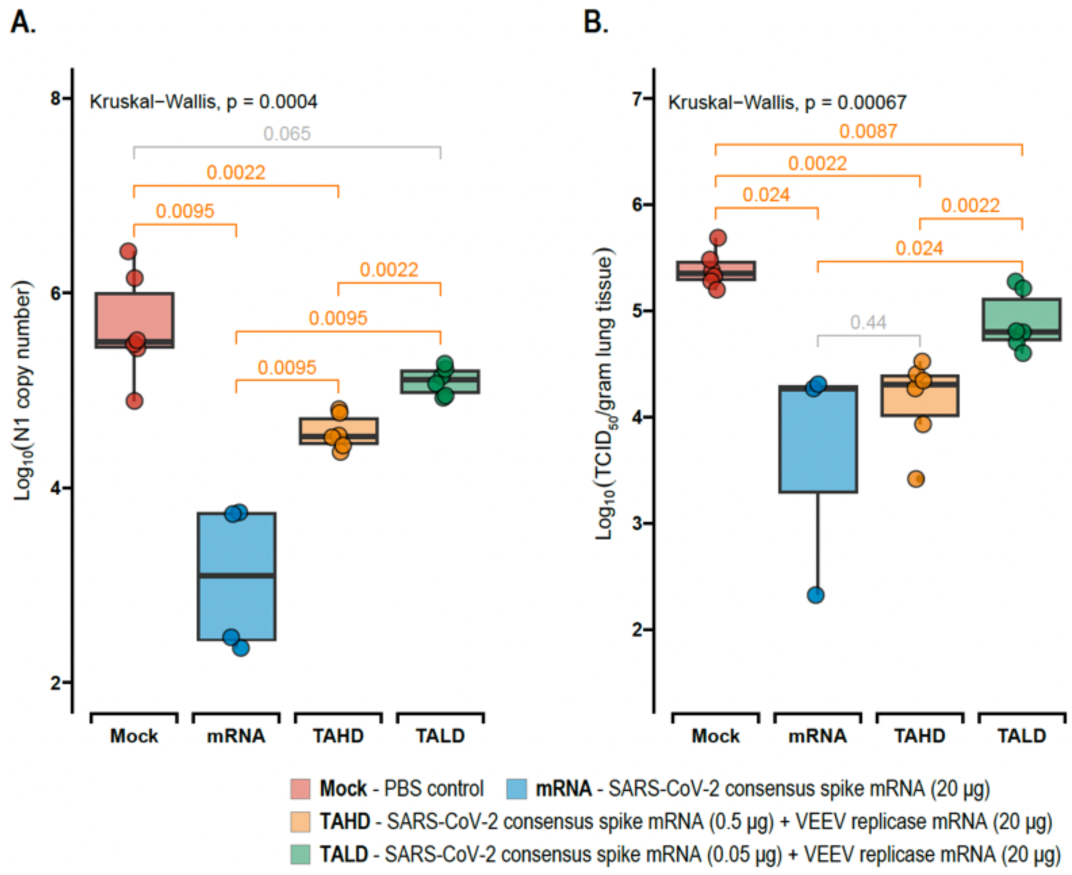
Assessment of viral load in mice following SARS-CoV-2 BA.1 challenge post-vaccination. Transgenic hACE2-expressing mice (*n*=6 per group; 3 males and 3 females) were vaccinated twice, two weeks apart, intra-dermally with either PBS (mock), SARS-CoV-2 consensus mRNA (mRNA), or two different doses of a trans-amplifying mRNA vaccine: high dose (TAHD) or low dose (TALD). Two weeks post-vaccination, mice were challenged intra-nasally with 10^4^ TCID50 of SARS-CoV-2 Omicron BA.1. Four days after the challenge, lung tissues were collected, homogenized, and analyzed for the presence of SARS-CoV-2 N1 viral gene by qPCR and for infectious virus titers by TCID50 assay. **(A)** qPCR results showed a significant difference in N1 copy numbers among the groups (p=0.0004, Kruskal-Wallis test followed by Wilcoxon Rank Test for pair-wise comparison). The mock group exhibited the highest N1 load, with significantly lower viral loads observed in the mRNA group (p=0.0095) and TAHD group (p=0.0022). No significant difference in viral load was found between the mock and TALD groups (p=0.065). The mRNA group had the lowest N1 levels compared to both the TAHD group (p=0.0095) and the TALD group (p=0.0095). Additionally, the TAHD group had lower N1 levels compared to the TALD group (p=0.022), indicating a dose-dependent response. However, the presence of the N1 gene does not necessarily indicate live virus presence, underscoring the importance of the TCID50 assay as a true measure of infectious viral load. **(B)** Infectious virus titers from homogenized lung samples showed a significant difference among the groups (p=0.00067, Kruskal-Wallis test followed by Wilcoxon Rank Test for pair-wise comparison). The mock group had the highest levels of infectious virus, significantly differing from all other groups: mRNA (p=0.034), TAHD (p=0.022), and TALD (p=0.0087). The mRNA group had infectious virus levels comparable to the TAHD group (p=0.44), demonstrating that the TAHD vaccine was able to reduce the challenge virus to levels similar to those achieved with the mRNA vaccine, despite having 40 times less antigen mRNA load. The mRNA group had lower infectious virus levels compared to the TALD group (p=0.024), and the TAHD group also had lower levels than the TALD group (p=0.0022). The box plots display individual values with median values (thick black line in the middle) and 95% confidence intervals.

Log_10_ median TCID_50_ titers per gram of lung tissue were calculated as 5.35 (95% CI: 5.21 to 5.57) in the mock-vaccinated group, 4.27 (95% CI: 0.81 to 6.45) in mice vaccinated with 20 μg of consensus spike mRNA (3 mice showed undetectable virus titers), 4.31 (95% CI: 3.72 to 4.58) in mice vaccinated with TA mRNA-high dose, and 4.80 (95% CI: 4.61 to 5.19) in mice receiving TA mRNA-low dose (**Fig. 4B**). Kruskal-Wallis test followed by Wilcoxon Rank Test for pair-wise comparison demonstrated significant reductions in TCID_50_ titers in mice vaccinated with consensus spike mRNA (p = 0.024), as well as those receiving either TA mRNA-high dose (p = 0.0022) or TA mRNA-low dose (p = 0.0087) compared to the mock-vaccinated group. There was no significant difference in TCID_50_ titers between mice vaccinated with consensus spike mRNA and those receiving TA mRNA-high dose (p = 0.44). However, the consensus spike mRNA group had significantly lower titers than the TA mRNA-low dose group (p = 0.024); And the TA mRNA high-dose group exhibited significantly lower titers compared to the TA mRNA-low dose group (p = 0.0022).

Overall, the TA mRNA-high dose vaccine resulted in a greater than 1 Log_10_ (>10-fold) reduction in lung viral titers, as measured by both qPCR (N1 gene copy number) and TCID_50_ assays, at day 4 post-challenge. This level of reduction was comparable to that observed in the consensus spike mRNA group, despite the TA mRNA-high dose vaccine containing 40 times less antigen mRNA (**Fig. 4A & 4B**).

### VEEV replicase suppresses anti-viral immune gene responses post-transfection in epithelial cells

Transcriptomic analysis of A549 cells transfected with VEEV replicase mRNA and SARS-CoV-2 mRNA (treatment group, n=3) versus cells transfected with unrelated mRNA and SARS-CoV-2 mRNA (control group, n=3) revealed significant alterations (p-value cut-off, p^0.05) in gene expression. At both 4- and 12-hours post-transfection, a notable downregulation (log_2_ fold change cut-off, ± 2.00) of interferon-stimulated genes (ISGs) and other immune-related genes, including IFNL2, IFNL3, IFNB1, CCL5, RAET1L, KRT17, and SEMA3D, was observed (**Fig. 5A & 5B**). Specifically, 28 genes were downregulated at 4 hours and 57 genes at 12 hours, with 20 genes consistently downregulated at both time points (**Fig. 5C**). Among these, key ISGs such as CCL5, IFNB1, IFNL1, IFNL2, IFNL3, and IFNL4 showed sustained suppression throughout the treatment (**Fig. 5D**). Gene Ontology (GO) term enrichment analysis indicated that immune response-related pathways were consistently suppressed at both 4- and 12-hours, with significant suppression observed in anti-viral pathways, including type I interferon signaling, serine phosphorylation of STAT proteins, and cytokine-mediated signaling at 12 hours (**Fig. 5E & 5F**).

**Fig. 5.**
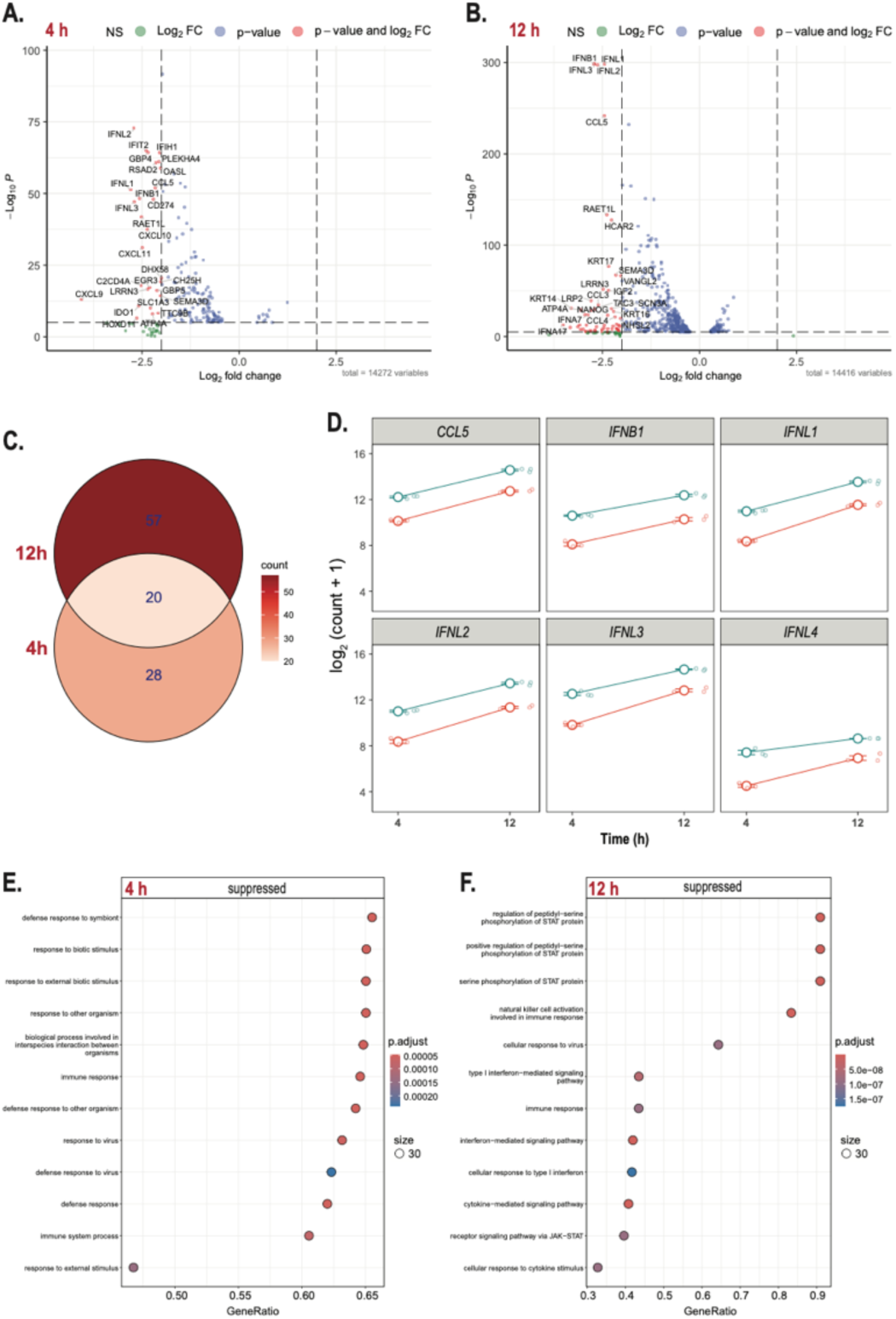
Transcriptomic analysis of A549 cells transfected with VEEV replicase mRNA and SARS-CoV-2 mRNA (treatment group, *n*=3) compared to cells transfected with an unrelated mRNA and SARS-CoV-2 mRNA (control group, *n*=3). (A &. **B)** Volcano plots showing differentially expressed genes at 4 hours **(A)** and 12 hours **(B)** post-transfection. Cut-off values of log2 fold change and p-value were set to ± 2.00 and 0.05. Significant downregulation of interferon-stimulated genes (ISGs) and other immune-related genes, such as IFNL2, IFNL3, IFNB1, CCL5, RAET1L, KRT17, and SEMA3D, were observed. **(C)** Venn diagram comparing the overlap of suppressed genes at 4 hours and 12 hours post-transfection, in the treatment group. A total of 20 genes were consistently downregulated at both time points, with 57 genes uniquely suppressed at 12 hours and 28 genes at 4 hours. **(D)** Combined dot and line plots demonstrating the expression of selected key ISGs (CCL5, IFNB1, IFNL1, IFNL2, IFNL3, IFNL4) over time, showing sustained suppression at 12 hours post-transfection in the cells transfected with the treatment group (indicated by red lines) compared to the control group (blue lines). **(E & F)** Gene Ontology (GO) term enrichment analysis of suppressed genes at 4 hours **(E)** and 12 hours **(F)**. Pathways related to immune response remained consistently suppressed at both time points. At 12 hours, significant suppression was observed in crucial pathways involved in anti-viral responses, including type I interferon signaling, serine phosphorylation of STAT proteins, and cytokine-mediated signaling. The size of the circles represents the gene ratio within each GO term, and the color gradient corresponds to the adjusted P-value. These findings highlight the temporal dynamics of immune response modulation following transfection of epithelial cells with the VEEV replicase mRNA and SARS-CoV-2 mRNA (treatment group), emphasizing the plausible role of VEEV replicase in the prolonged suppression of critical antiviral pathways at later time points.

## Discussion

The continuous emergence of novel SARS-CoV-2 variants poses significant challenges to the sustained efficacy of current vaccines. The available vaccines were updated in 2022 and 2023. However, by the time these vaccines were manufactured and deployed, a different variant had become predominant, reducing their efficacy in neutralizing the circulating SARS-CoV-2 strains^38^. We designed and evaluated a trans-amplifying (TA) mRNA vaccine to elicit neutralizing antibodies against multiple variants of the virus while concurrently minimizing the dosage requirement of the antigenic mRNA. Immunization of mice with the TA mRNA vaccine induced broadly cross-neutralizing antibody response against SARS-CoV-2 B.1 lineage and multiple variants, Alpha, Beta, Gamma, Delta, and Omicron BA.1. Mice vaccinated with the TA mRNA vaccine had a greater than 10-fold reduction in viral load following challenge with SARS-CoV-2. The TA mRNA vaccine with 0.5 µg of consensus spike mRNA elicited antibody responses comparable to those of a conventional mRNA vaccine formulated with 20 µg of consensus spike mRNA (40 times higher than the TA mRNA dose), demonstrating a significant dose-sparing effect.

The breadth of neutralizing antibodies is a critical determinant of SARS-CoV-2 vaccine efficacy ^39, 40^. Consensus spike protein antigen had four amino acid substitutions (K417N, T478K, E484A, N501Y) within the receptor binding domain (RBD) portion and two outside the RBD (D614G and H655Y), as compared to the ancestral SARS-CoV-2 spike sequence. Of these, K417N and N501Y substitutions were earlier known to stabilize the spike protein by decreasing the flexibility and promoting open conformation of the RBD ^41, 42, 43, 44^. D614G and H655Y are convergent adaptations to the polybasic S1/S2 cleavage site and were shown to enhance the stability of the S1 subunit in the open RBD conformation ^45, 46, 47, 48, 49^. The open conformation of RBD exposes the highly conserved epitopes in the spike protein, potentially increasing the immunogenicity of the spike protein ^50^. Additionally, we introduced mutations previously employed to stabilize the spike protein in its prefusion conformation, including the GSAS substitution at residues 682-685 and six proline mutations at positions F817P, A892P, A899P, A942P, K986P, and V987P ^37, 46^. These six proline substitutions were earlier proven to enhance the stability of the spike protein ^37^. In this study, mice vaccinated with consensus spike mRNA alone or as a TA mRNA vaccine resulted in the generation of neutralizing antibodies against multiple SARS-CoV-2 variants. Consistent with our findings, a separate study that evaluated the efficacy of a spike-protein vaccine developed through homology of SARS-CoV-2 virus variants demonstrated protection against multiple virus variants ^51^. A similar strategy utilizing consensus proteins for vaccine design was tested successfully with Avian Coronavirus, MERS-CoV, Influenza, and Dengue ^34, 35, 36, 52, 53, 54^. Hence, a consensus spike protein strategy offers a promising approach for developing a universal SARS-CoV-2 vaccine, potentially eliminating the need for frequent updates and ensuring broader, long-term protection against emerging variants.

mRNA vaccines require a considerably high dose of antigenic mRNA to elicit the necessary immunogenic responses (Moderna’s Spikevax® contained 100 µg/dose, while Pfizer/BioNTech’s Comirnaty® contained 30 µg/dose)^55^. Significant gaps exist in both mRNA demand and supply, especially in developing and underdeveloped countries. Amplifying mRNA vaccines represent an advancement over conventional mRNA vaccines, particularly as TA mRNA vaccines provide opportunities for a reduced requirement of antigenic mRNA and enhanced modularity. TA mRNA vaccines using replicase from Semliki forest virus (SFV) and Chikungunya virus were investigated for influenza, Chikungunya, and Ross River viruses ^23, 29, 56^. These studies demonstrated the capacity of TA mRNA vaccines to decrease the antigenic mRNA dose in the tested formulations ^23, 29, 56^. Our vaccine was developed based on the comprehensive characterization of VEEV replicase activity ^57, 58^. The VEEV replicase was flanked by the 5’ untranslated region (UTR) of the human alpha-globin gene and the 3’ UTRs of AES and mtRNR1, enhancing translation efficiency and RNA stability ^23, 59^. Further, this design also ensures that the VEEV replicase is not amplifiable by the replicase itself, thereby enhancing the safety of the constructs. The identification of antigenic mRNAs by the VEEV replicase was accomplished by flanking conserved sequence elements of the VEEV ^57, 58^. We demonstrated through *in vitro* experiments that cells transfected with TA mRNA constructs exhibited an increase in antigenic protein expression over time. In mice, the TA mRNA vaccine allowed a 40-fold conservation of antigenic mRNA dose compared to a conventional mRNA vaccine.

Transcriptomic analysis of A549 cells transfected with TA mRNA vaccine constructs revealed sustained suppression of key antiviral genes, providing novel insights into the potential immunomodulatory effects of the TA mRNA vaccine platform. Specifically, the downregulation of immune-related genes, particularly interferon-stimulated genes such as CCL5, IFNB1, IFNL1, IFNL2, IFNL3, and IFNL4, suggests prolonged modulation of the host immune response by the VEEV replicase. VEEV nonstructural protein 2 (nsp2), a key component of the viral replicase, is known to inhibit host protein translation and suppress the antiviral response by downregulating interferon signaling pathways ^60^. The observed suppression of type I interferon signaling, serine phosphorylation of STAT proteins, and cytokine-mediated signaling at 12 hours suggests potential interference with the JAK-STAT pathway. This is consistent with previous findings that VEEV replicons can impair STAT1 phosphorylation and nuclear translocation^61^. Additionally, reduced expression of IL-6 in the VEEV replicase mRNA group compared to the control (**Supplementary data.5**) may have contributed to the observed suppression of STAT protein phosphorylation. The observed suppression of antiviral pathways, particularly type I interferon signaling, may enhance the efficacy of the TA mRNA vaccine by creating a more favorable environment for mRNA translation and antigen production. This modulation potentially allows for increased expression of the SARS-CoV-2 spike protein, contributing to a stronger immune response.

This study has several limitations that offer important insights for guiding future research. Nucleoside modifications are essential components of the contemporary mRNA vaccines, which mitigate post-vaccination inflammatory responses ^62^. The mRNA constructs utilized in the present study did not include nucleoside modifications, necessitating an assessment of their impact on the vaccine’s efficacy. The capacity of the consensus SARS-CoV-2 spike protein to neutralize variants that have emerged subsequent to Omicron BA.1 remains to be experimentally validated. The immunogenicity of the VEEV replicase requires additional inquiry, as it is plausible that antibodies elicited against the replicase may influence the effectiveness of forthcoming booster doses. Furthermore, VEEV replicase is noted to suppress antiviral pathways, such as type I interferon signaling, which necessitates further investigation into the safety profile of the vaccine. Future research efforts should also explore the potential benefits of multiplexing TA mRNA vaccines, whereby mRNAs from multiple SARS-CoV-2 variants are combined to enhance immune responses. In addition, nanoparticle-based delivery systems could significantly improve immunogenicity and overall vaccine efficacy.

In conclusion, our investigation of the TA mRNA vaccine underscores its potential as an effective and adaptable strategy to address the challenges posed by SARS-CoV-2 and other highly variable RNA viruses with pandemic potential. The TA mRNA vaccine employing a consensus spike demonstrated robust immunogenicity, eliciting cross-neutralizing antibody responses against a broad spectrum of SARS-CoV-2 variants, including major variants of concern. A key advantage of this platform is its significant dose-sparing capability, achieving comparable immune responses with 40 times less antigen mRNA than the conventional spike mRNA vaccine. Further optimization of this TA mRNA platform with a consensus spike protein, could be valuable in the ongoing fight against SARS-CoV-2 and its evolving variants. Additionally, the insights gained from this approach provide a strong foundation for developing universal coronavirus vaccines and broader strategies for combating future pandemic-prone RNA viruses with high genetic variability.

## Methods

### Plasmids for consensus spike and replicase

The VEEV replicase sequence was sourced from plasmid pTK194, kindly provided by Ron Weiss (Addgene plasmid #59563; http://n2t.net/addgene:59563; RRID)^63^. This sequence was codon-optimized for expression in human cells using the GenSmart™ Codon Optimization Tool (Genscript, USA). The optimized sequence was flanked by the 5’ UTR of the human alpha-globin gene and the 3’ UTRs of AES and mtRNR1^23, 55, 59^. Following the 3’ UTR, a polyadenylation (Poly A) tail was included, with SapI restriction sites positioned downstream to ensure a precise and clean Poly A tail ^64^. Upstream of the 5’ UTR, an AscI restriction site and a T7 promoter sequence were introduced (**Fig. 1D**). The sequence is provided in **Supplementary data.1.**

The spike protein sequence was generated by constructing a consensus of spike protein sequences from multiple SARS-CoV-2 variants (GenBank accession numbers: QHD43416.1 [Ancestral B.1], QWE88920.1 [Alpha], QRN78347.1 [Beta], QRX39425.1 [Gamma], QUD52764.1 [Delta], UFO69279.1 [Omicron BA.1/BA.2], UMZ92892.1 [Omicron BA.2.12.1], UPP14409.1 [Omicron BA.4], UOZ45804.1 [Omicron BA.5]) (**Fig. 1A**). The alignment was conducted using MUSCLE within SnapGene software (version 7.0.3). To stabilize the pre-fusion state of the spike protein, six proline substitutions were incorporated, following the method described by Hsieh et al. ^37^. The consensus spike sequence was flanked by conserved VEEV replicase sequence elements. As with the replicase, an AscI restriction site and a T7 promoter sequence were placed upstream of the 5’ UTR, and a Poly A tail with a SapI restriction site followed the 3’ UTR (**Fig. 1D**). The spike protein sequence was optimized for expression in human cells using the GenSmart™ Codon Optimization Tool (Genscript, USA). The sequence is provided in **Supplementary data.3.** Structural modeling confirmed the stability of the pre-fusion conformation of the synthesized consensus spike protein (**Fig. 1B**).

Both the replicase and spike protein sequences were synthesized in pBluescript II SK(+) plasmids with ampicillin resistance (Genscript, USA). The plasmids were subsequently transformed into DH5α E. coli for large-scale plasmid production.

### Synthesis of mRNA from plasmid DNA

Plasmids were linearized by restriction digestion with AscI and SapI. The linearized plasmids were then transcribed *in vitro* utilizing the MEGAscript™ T7 Transcription Kit (Cat.# AM1334) from Invitrogen, Thermo Fisher, USA. The resulting RNA was capped using the Vaccinia Capping System (Cat.# M2080S) and mRNA Cap 2’-O-Methyltransferase (Cat.# M0366S) from NEB, USA. The capped RNA was subsequently purified with the Monarch® RNA Cleanup Kit (Cat.# T2040L) from NEB, USA. The quality and quantity of the resulting mRNA were analyzed and confirmed using the 4200 TapeStation System from Agilent, USA.

### Transfection of mRNA in cell culture

To evaluate the protein expression of the designed mRNAs, A549 cells (Human Lung Carcinoma Cells, Cat.# CCL-185, ATCC, USA) were transfected with 2.5 μg of VEEV replicase mRNA and 0.05 μg of SARS-CoV-2 mRNA (accompanied by 2.5 μg of unrelated or scrambled mRNA) using Lipofectamine™ MessengerMAX™ Transfection Reagent (Cat.# LMRNA015, Thermo Fisher Scientific, USA). Cells were transfected with either mRNA alone or in combination, and subsequent analyses were conducted at 4 and 12 hrs post-transfection.

### Western blot analysis of consensus spike and replicase expression in transfected cells

At given time-points post-mRNA transfection, total protein lysate was collected using RIPA Lysis and Extraction Buffer (Cat.# 89900) from Thermo Fisher Scientific, USA. The cell lysates were separated on 10% polyacrylamide pre-cast gels (Cat.# 4561035), Biorad, USA, and subsequently transferred onto a polyvinylidene difluoride (PVDF) membrane. The membrane was probed with SARS-CoV-2 Spike Protein S1/S2 Polyclonal Antibody (Cat.# PA5-112048), Thermo Fisher Scientific, USA, or Non-structural polyprotein Antibody against Venezuelan equine encephalitis virus (strain Trinidad donkey) (VEEV) Non-structural polyprotein (Cat.# CSB-PA329710XA41VAZ), Cusabio, USA. Following that, the membrane was treated with Goat Anti-Rabbit IgG Antibody, Fc, HRP conjugate (Cat.# AP156P), Sigma-Aldrich, Millipore Sigma, USA. Additionally, the lysates were probed with Anti-β-Actin−Peroxidase antibody, Mouse monoclonal (Cat.# A3854), Sigma-Aldrich, Millipore Sigma, USA. The blots were visualized for chemiluminescence using SuperSignal™ West Femto Maximum Sensitivity Substrate (Cat.# 34094), Thermo Fisher Scientific, USA.

### Immunogenicity studies of trans-amplifying mRNA vaccine in mice

K18-hACE2 transgenic mice aged between five to seven weeks were procured from The Jackson Laboratory, USA (Strain #034860; B6.Cg-Tg(K18-ACE2)2Prlmn/J)^65^. The mice were organized into four groups, each consisting of six individuals (3 males and 3 females). The groups were designated as follows: Mock (received phosphate buffered saline), Consensus Spike mRNA (20 μg/dose), TA mRNA-high dose: Consensus Spike mRNA (0.5 μg/dose) + Replicase mRNA (20 μg/dose), and TA mRNA-low dose: Consensus Spike mRNA (0.05 μg/dose) + Replicase mRNA (20 μg/dose).

Immunizations were administered intra-dermally (50 μL/mouse) under isoflurane anesthesia, with two doses given two weeks apart. Blood was collected on day 28 for serum analysis. On day 30, all mice were intranasally challenged with 10^4 TCID_50_ (20 μL/mouse) of SARS-CoV-2 Omicron BA.1: Isolate hCoV-19/USA/MD-HP20874/2021 (Lineage B.1.1.529; Omicron Variant) from NR-56461, BEI Resources, USA. Four days post-challenge, the mice were euthanized with CO_2_, and lung tissues were collected. A portion of the lung was frozen for infectious virus load titration, while another part was stored in RNA later. Both sections underwent homogenization, and subsequent analyses were performed. The experimental plan and the details are given in **Fig. 2A & 2B.**

### Virus neutralization assay

The serum samples’ virus-neutralizing titers were assessed using a cell-based assay with SARS-Related Coronavirus 2 (SARS-CoV-2), Isolate hCoV-19/USA/PHC658/2021 (Lineage B.1.617.2; Delta Variant) from NR-55611, BEI Resources, USA^66^. Monolayers of Vero C1008 cells (Vero 76, clone E6, Vero E6 cells), CRL-1586, ATCC, USA were cultivated in 96-well microtiter plates. Heat-inactivated serum samples were diluted two-fold in triplicate and incubated with 100 TCID_50_ virus in a CO_2_ incubator at 37°C for 1 hr. The serum–virus mixture was then added to cell monolayers and further incubated for 3 days in a CO_2_ incubator at 37°C. Plates were observed for cytopathic effects such as rounding and sloughing off cells. The reciprocal of the highest serum dilution where at least two of the three wells demonstrated protection (no cytopathic effect) was determined as the neutralizing titer of the sample.

### Pseudovirus neutralization assay

The sera from the study were subjected to a SARS-CoV-2 pseudovirus neutralization assay using 293T-hACE2 cells, which are Human Embryonic Kidney Cells expressing Human Angiotensin-Converting Enzyme 2 (HEK-293T-hACE2 Cell Line, NR-52511, BEI Resources, USA), following the previously described procedure^67^. SARS-CoV-2 spike pseudoviruses were generated using lentiviral packaging plasmids (SARS-Related Coronavirus 2, Wuhan-Hu-1 Spike-Pseudotyped Lentiviral Kit V2 (Plasmid/Vectors), NR-53816, BEI Resources, USA). Plasmids encoding spike proteins of SARS-CoV-2 variants were procured from Addgene, USA (Alpha (Cat.# 170451), Beta (Cat.# 170449), Gamma (Cat.# 170450), Delta (Cat.# 172320), Omicron BA.1 (Cat.#180375), Omicron XBB.1 (Cat.#194494)^68, 69, 70, 71^. Duplicate two-fold serial dilutions of serum samples were incubated with 100 µL of pseudoviruses (equivalent to 10^4 -10^5 relative luminescence units (RLU)) at 37 °C for 1 hour in a CO_2_ incubator. Subsequently, 13,000 293T-hACE2 cells per well were added and incubated at 37 °C for 2 days in a CO_2_ incubator. After 2 days, 100 µL/well of bright-glo luciferase reagent (Promega, Cat. No. E2620) was added to the wells, and luminescence was measured. The RLUs were recorded, and a half-maximal effective concentration titer (EC_50_) for each serum sample was calculated from dose-response curves using a four-parameter Hill equation using R package/version ‘*drc’* (version 3.0-1)^72^.

### Estimation of viral load in mice post SARS-CoV-2 challenge

The lung tissue, preserved in RNA later, was homogenized with DPBS and subsequent RNA extraction using the MagMAX™-96 Total RNA Isolation Kit (Cat.# AM1830) from Applied Biosystems™, Thermo Fisher Scientific, USA. Complementary DNA (cDNA) from the extracted RNA was prepared using qScript cDNA SuperMix (Cat.# 95048-100) from Quantabio, USA. Results were normalized with the Eukaryotic 18S rRNA Endogenous Control (FAM™/MGB probe, non-primer limited) (Cat.# 4333760F) from Applied Biosystems, Thermo Fisher Scientific, USA ^73^. The cDNA was then analyzed for SARS-CoV-2 N1 by qPCR. The primers and probe were sourced from IDT, USA (Cat.# 10006713) ^74^. The master mix utilized for qPCR reactions was TaqMan™ Fast Universal PCR Master Mix (2X), no AmpErase™ UNG (Cat.# 4364103), Applied Biosystems, Thermo Fisher Scientific, USA.

Another portion of the lung from each mouse was homogenized, and the clarified supernatant was assessed for infectious virus titers. Virus yield was determined through an endpoint dilution assay on VeroE6 cells that calculates the 50% tissue culture infective dose (TCID_50_) using the Reed and Muench method ^75^.

### Transcriptome analysis

RNA was extracted from cell cultures of A549 after 4 hours and 12 hours of transfection with mRNAs encoding 2.5 μg of VEEV replicase mRNA and 0.05 μg of SARS-CoV-2 mRNA. Another set of cells was similarly transfected with 2.5 μg of an unrelated mRNA and 0.05 μg of SARS-CoV-2 mRNA, and RNA was extracted from these cells. The RNeasy Plus Mini Kit (Cat.# 74134) from Qiagen, Germany was used for RNA extraction, while the NEBNext® Poly(A) mRNA Magnetic Isolation Module (Cat.# E7490L) from NEB, USA was used to enrich mRNA from the extracted RNA. RNA libraries were then prepared using the xGenTM RNA Lib Prep 96rxn xGen RNA Library Prep Kit (Cat.# 10009814) from IDT, USA, and normalized using the xGenTM Normalase^TM^ Module (Cat.# 10009793) from IDT, USA. The Illumina NextSeq 2000 was used to sequence the prepared libraries. The quality of reads was checked via FASTQC, and then mapped to the human reference genome (GRCh38.p14) using HISAT2 with default parameters ^76, 77, 78^. The mapped reads were counted using featureCounts, and the results were analyzed via DESeq2 using the default Benjamini-Hochberg correction of the *p*-values from the Wald test ^79, 80^. A 0.05 False Discovery Rate (FDR) cut off was considered to classify genes as differentially expressed. The clusterProfiler R package was used for gene ontology enrichment analysis using the biological process ontology ^81^.

### Statistical analysis

Statistical significance between groups was calculated using Kruskal-Wallis test followed by Wilcoxon Rank test for pair-wise comparison in the R package/version ‘*ggpubr*’ package (version 0.6.0)^82^. A *P* value of less than 0.05 was considered statistically significant.

## Ethical statement

All experiments were conducted following the necessary approvals from the Institutional Biosafety Committee (BIO202300052, BIO202300086) and Institutional Animal Care and Use Committee (PROTO202001704) at The Pennsylvania State University, University Park, PA, USA.

## Data availability

All genomic sequences used for transcriptome analysis in this study are publicly available at https://www.ncbi.nlm.nih.gov/geo/query/acc.cgi?acc=GSE277947.

## Acknowledgments

The study was funded by the chair funds from the Huck Institutes of the life sciences, (SVK), and Interdisciplinary Innovation Fellowship at the One Health Microbiome Center (AG), Pennsylvania State University. The authors express their sincere gratitude to the Animal resource program (Sima Bruggeman, Jeffery Dodds, and Melissa Welker), Pell staff (Lindsey Combs and Jacob Perryman), and Lingling Li for their assistance in conducting experiments. We also appreciate the technical assistance provided by Dr. Neela Yennawar and Dr. Rajeswaran Mani from the Huck Core Facilities at Pennsylvania State University. We acknowledge the use of GraphPad PRISM, R, Grammarly, and ChatGPT for data analysis and manuscript preparation.

## Author contributions

Project concept (A.G., S.V.K.); experimental design (A.G. S.V.K., M.A); acquired data (A.G., R.H.N., S.K.C., S.R., P.J., M.B., L.C.L., M.S.N.); analyzed data (A.G., S.V.K., S.M.); prepared figures (S.M., A.G.); Prepared manuscript (A.G., S.M., S.V.K); Supervision and funding acquisition (S.V.K). All authors reviewed the manuscript and gave final approval for publication.

## Supplementary data

1. **DNA sequence of linearized VEEV replicase** cgcgcctaatacgactcactatagggagaataaattcttctggtccccacagactcagagagaacccgccaccatgggcggcgcatgagagaagcccagaccaattacctacccaaaatggaaaaggtgcacgtggacatcgaggaagacagcccctttctgcgggccttgcagagaagcttccctcagttcgaggtggaagccaagcaggtcactgataatgaccatgctaatgccagagcgttttcgcatctggcctctaaactgatcgagacagaggtggatccctctgatacaatcctggacatcggcagcgcccctgctagacggatgtacagcaagcacaagtaccactgcatctgccctatgcggtgcgccgaagatccagatcggctgtacaagtacgccaccaagctgaaaaagaactgcaaggagatcaccgataaggagctggacaagaagatgaaggagctggccgctgtgatgagcgacccagacctggaaacagagacaatgtgcctgcacgacgacgagagctgcagatacgagggacaggtagccgtgtaccaggacgtgtacgctgtggatggccctaccagcctgtaccaccaggccaacaagggcgtgcgggtggcctactggatcggatttgacaccacacctttcatgttcaagaacctggccggcgcctatcctagctactccaccaactgggccgacgagacagtgctgacagccagaaacatcggcctgtgtagctctgatgtgatggaaagaagcagacggggcatgtctatcctgagaaagaaatatctcaagcctagcaacaacgtgctgttcagcgtgggcagcacaatctaccacgagaagcgggacctgctgagatcttggcaccttcctagcgtcttccatctgagaggaaagcagaactacacctgtagatgtgaaaccatcgtgtcttgtgacggctacgtggtgaagagaatcgccatctcccctggactctacggcaaaccttcaggttatgccgctacaatgcaccgggaaggcttcctgtgttgcaaggtgaccgacaccctgaacggcgagagagtgtccttccccgtgtgcacctacgtgcctgccactctgtgcgaccaaatgaccggcatcctggctacagacgtctccgccgatgatgcccagaagctgctggtgggcctgaaccagcgcatcgtggtgaatggccggacacagcggaacaccaacaccatgaagaattacctgctgcctgtggttgcccaggctttcgccagatgggcgaaggaatacaaggaggaccaggaggatgaaagacctctgggcctgcgggacagacagctggtcatgggctgctgctgggccttcagaaggcataagattaccagcatctacaagcggcctgatacccaaaccatcatcaaggttaacagcgacttccacagcttcgtgctgcctagaattggcagcaataccctggaaatcggcctgaggaccagaatcagaaagatgctggaggagcacaaagaacccagcccactgatcaccgccgaggacgtgcaggaggccaagtgcgctgcagacgaagccaaagaggtcagagaggccgaggaactgcgggccgccctgccccccctggctgctgatgtggaggagcccaccctggaggccgacgtggacctgatgctgcaagaggccggagccggatccgttgagacacctagaggcctgatcaaggtcaccagctatgacggcgaagataagatcggctcttacgccgtgctgtctcctcaagctgtgctgaagtcagaaaagctgagctgtatccaccctctggctgagcaggtgatcgtgatcacccactctggacggaaaggaagatacgccgtagagccttaccacggcaaggtggtggtgccagagggccacgccatccccgtgcaggacttccaagccctgagcgagtctgccacgatcgtgtacaacgagcgggaattcgtgaaccggtacctgcaccacatcgccacccatggaggcgccctcaatacagatgaagaatactacaagaccgtgaagccatccgaacacgacggcgagtacctgtacgatatcgaccggaagcaatgcgtgaaaaaagaactggtgacaggtctgggcctgaccggcgagctggtcgacccacccttccacgagttcgcctacgagagcctgagaaccagacccgccgccccttatcaggtgcccacgattggagtatacggcgtgcctgggagcggcaagagcggcatcatcaaaagcgccgtgaccaagaaggacctggtcgtgagcgccaagaaagaaaactgcgccgagatcatccgggacgtgaagaaaatgaaaggccttgacgtgaacgctagaaccgtggacagcgtgctgctgaatggctgcaagcaccccgtggagaccctgtacatcgacgaggccttcgcctgccacgccggcaccctgagagctctgatcgccatcatcagacctaagaaggccgtgctgtgcggcgatcctaagcagtgtggctttttcaacatgatgtgcctgaaggtgcacttcaaccacgaaatctgcacccaggtctttcacaagagcatcagcagacggtgtaccaaatctgtgacctccgtggtctctacactgttctacgacaagaaaatgcgtacaaccaaccccaaggagacaaagatcgttatcgacaccaccggcagcaccaagcccaagcaagacgatctgatcctgacctgcttccgcggctgggtgaagcagctgcagatcgattacaagggcaatgaaatcatgaccgccgctgccagccagggcctgacaagaaagggcgtttacgccgtgcggtacaaggtgaacgaaaaccccctgtacgcccctaccagcgagcacgtgaatgtcctcctgaccagaactgaggaccgcatagtgtggaagacactggccggagacccctggatcaagaccctgacagctaagtaccctggcaacttcaccgccaccatcgaggaatggcaggccgagcacgacgccatcatgcggcacatcctggaacggcctgatcctacagatgtgttccagaacaaggctaatgtgtgctgggccaaagccctggtgccggtgctgaaaaccgccggcattgatatgaccacagagcagtggaacaccgtggactacttcgagaccgacaaggcccacagcgccgagatcgtactgaaccagctgtgcgtgagattcttcggcctggacctggacagcggcctgttttctgcccctaccgttccactgagcatcagaaacaaccattgggacaacagcccttcacctaacatgtacggcttaaacaaggaagtggtgaggcagctgagccggagatacccccagctgcctagagctgtggccaccggaagagtgtacgatatgaacactggaaccctgagaaactacgaccctcggatcaacctggtgcctgtgaatagaagactccctcacgccctggtgctgcatcacaacgagcaccctcagagcgacttctcttcttttgtgagcaagctgaagggcagaacagtgctcgtggtgggcgagaagctgagcgtgccaggcaagatggtggactggctgagcgacagacctgaagctacctttcgggccagactggatctgggcattcccggagatgtgccaaaatacgacatcatcttcgtgaacgtgcggacaccttacaaataccaccactaccagcagtgcgaggatcacgccatcaagctgagcatgctgacaaagaaggcctgtctgcacctgaaccccggcggcacatgtgtgagcatcggatatggctatgctgatagagcctccgagagcatcatcggcgccatcgcaagactgttcaagttcagccgggtgtgcaaacccaagtccagcctggaagaaaccgaagtgctttt tgtgttcatcggctacgacagaaaggccagaacacataatccttacaagctgtcctccaccttaacaaacatctacaccggcagcagactgcacgaggccggatgtgctcctagctatcacgtggtgcgcggcgacatcgccaccgctaccgagggcgtgatcatcaacgccgccaatagcaagggccagcctggcggcggagtttgtggcgctctgtacaagaaattcccggagagcttcgacctgcaacctattgaggtgggcaaggcccggctggtgaagggcgccgcaaagcacatcattcatgctgttggccccaacttcaacaaggtgtccgaggtggaaggagataagcagctggccgaggcctacgagagcatcgccaaaatcgtgaacgacaacaattacaaaagcgtggccatccctctgctgagcaccggcatcttcagcggcaacaaagatagactcacccagagcctgaaccacctgctgaccgccctggacaccaccgacgccgacgtggctatctattgcagagataagaagtgggagatgaccctgaaggaagccgtggcccggagagaggctgtggaagaaatctgcatcagcgacgactctagtgtgaccgagcctgacgccgaactcgtgcgggtgcaccctaagtccagcctcgcgggcagaaagggctacagcaccagcgacggcaaaacattcagctacctggaaggcacaaagttccaccaggccgccaaggacatcgccgagatcaatgccatgtggcctgtggccacagaagctaacgagcaagtatgtatgtacatcctgggcgagagcatgagcagcatcaggtccaagtgccccgtggaagagagcgaggcgtctacccctccatccaccctgccttgcctgtgcatccatgccatgacaccagagagagtgcagagactgaaagcctctagacctgaacagatcacagtgtgttcctcttttcctctgcccaagtacagaatcacaggcgtgcagaagatccagtgcagccagcccatcctgtttagccctaaggtgcctgcttacatccaccccagaaagtacctggtcgagacacctcctgtggacgagacccctgaacctagcgccgaaaaccagtctaccgagggaaccccagaacagcctccactgatcaccgaggatgagacaagaacccggacacctgagcccatcattatcgaggaggaagaggaagatagcatcagcctgctgtctgacggccctacccaccaggtgctgcaggtggaggccgatatccacggccctccttctgtgagcagcagctcatggagcatcccccacgccagcgacttcgacgtggacagcctgagcatcctggatacactggaaggagccagcgtgacaagcggagccacatctgccgagactaactcttacttcgccaagagcatggaattcctggctagacctgtgcccgccccccggaccgtctttagaaaccccccccaccctgccccaagaaccagaacccctagcctggccccttctcgggcatgcagcagaaccagtctggtgagcacaccacctggtgtcaatagagtgatcacccgggaggagctcgaagccctgaccccatctcggaccccgtcacgatccgtgagccggaccagcctggtgtccaacccccccggcgtgaacagagtgatcacaagagaggaattcgaggccttcgtagcccagcaacagtgacggtttgatgcgggtgcatatatcttcagcagcgacacaggacagggccacctgcagcagaaaagcgtgcggcagaccgtgctgagcgaggtggtgctggaaagaacagaactggaaatctcttacgcccctagactggaccaggagaaggaagaattgctgcggaagaaactgcagctgaaccctacacctgccaaccggagcagataccagtcccggaaggtcgagaacatgaaggccatcaccgctaggagaatcctgcaaggcctgggccactacctgaaggctgagggcaaggtggagtgttacagaaccctgcaccccgtgcctctgtacagctcctccgtgaatagagccttttctagccccaaggttgccgtcgaagcctgcaacgccatgctgaaggagaacttcccaacagtggcctcttactgcatcatccccgagtacgacgcctacctggacatggtggacggcgcttcatgctgcctggacaccgcctccttctgtcctgctaagctgcggagcttccccaaaaagcacagctacctcgagcccaccatccggtctgctgtgccaagcgccatccagaacacactgcaaaacgtcctggccgccgcaaccaaacggaactgcaatgtgacccagatgagagaactgcctgtgctggatagcgccgcctttaacgtggaatgcttcaagaagtacgcctgcaacaacgagtactgggagacattcaaggaaaaccccatcagactgactgaggaaaatgtggtgaactacatcaccaagctcaagggccctaaggccgctgctctgttcgccaagacccacaacctgaacatgctgcaggacatccctatggacagattcgtgatggacctgaagagagatgtgaaggtgacccctggaaccaagcacacagaggagcggcctaaggtgcaggtgattcaggccgccgatcctctggctaccgcctatctctgtggcatccacagagagctggtgcggcggctgaatgccgtgctgctgcccaacatccacaccctgttcgacatgagcgccgaggacttcgatgctatcattgccgagcacttccagcctggcgactgcgtgctggaaaccgacatcgccagcttcgacaagtctgaggatgatgccatggccctaaccgccctgatgatcctcgaggacctgggcgtggacgccgaactgctgaccctgatcgaggccgctttcggcgagatcagcagcatccacctccctaccaagaccaagttcaagttcggcgccatgatgaaaagcggcatgttcctgacactgtttgtcaacaccgtgatcaacatcgtgatcgccagcagagtgctgagagagagactgaccggctcaccttgcgccgccttcatcggcgacgacaacatcgtgaaaggcgtgaagtctgataagctgatggctgatagatgcgccacctggctgaatatggaagtgaagatcatcgacgcagttgtgggagaaaaggctccatacttctgcggcggcttcatcctgtgcgacagcgtgaccggtacagcctgtagagtggccgaccccctgaaaagactgtttaagctgggaaaacctctggccgccgacgatgagcatgatgacgatcggagacgcgccctgcacgaggaatccaccagatggaacagagtgggaatcctgagcgaactgtgtaaagccgtggaaagcagatacgagacagtgggcacatctattatcgtgatggccatgacaacactggctagcagcgtcaagagctttagttacctgaggggcgcccctatcactctctacggctaacctgaatggacaacgttcctgcaggtacgacatagtctagtccgccaagtatacagcagcaattggcaagctgcttacatagaactcgcggcgattggcatgccgccttaaaatttttattttatttttcttttcttttccgaatcggctggtactgcatgcacgcaatgctagctgcccctttcccgtcctgggtaccccgagtctcccccgacctcgggtcccaggtatgctcccacctccacctgccccactcaccacctctgctagttccagacacctcccaagcacgcagcaatgcagctcaaaacgcttagcctagccacacccccacgggaaacagcagtgattaacctttagcaataaacgaaagtttaactaagctatactaaccccagggttggtcaatttcgtgccagccaccgcattttgtttttaatatttcaaaaaaaaaaaaaaaaaaaaaaaaaaaaaaaaaaaaaaaaaaaaaaaaaaaaaaaaaaaaaaaaaaaaaaaaaaaaaaaaaaaaaaaaaaaaaaaa

**2. Validation of TA mRNA vaccine design using nano-luciferase as the gene of interest**

**Fig.**
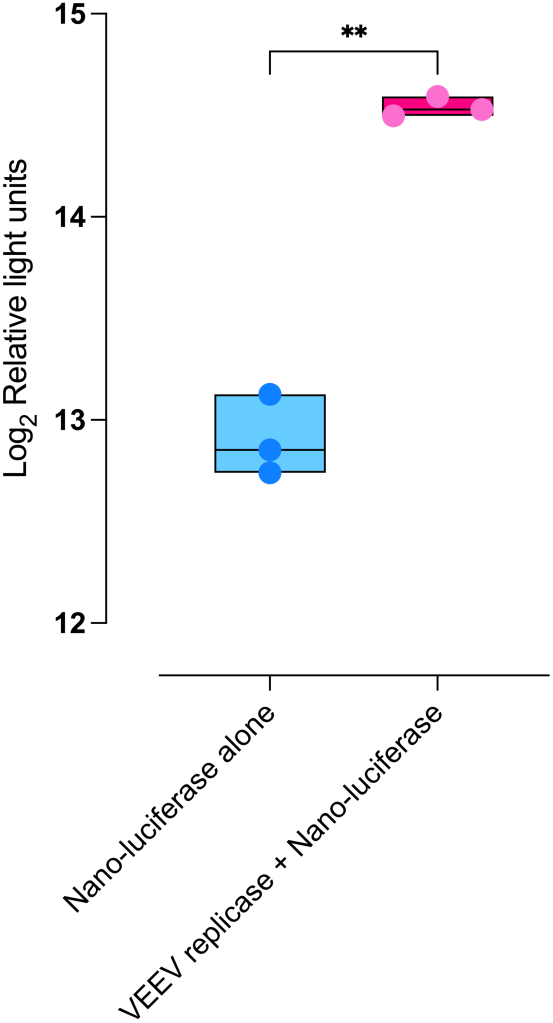
Luciferase expression in 293T cells transfected with mRNA encoding nano-luciferase alone (control, *n*=3) and cells mRNA co-transfected with mRNAs encoding VEEV replicase and nano-luciferase (treatment, *n*=3). The treatment group exhibited significantly higher luciferase expression compared to the control (p = 0.0033, t-test).

3. **DNA sequence of linearized SARS-CoV-2 consensus spike** cgcgcctaatacgactcactatagggatgggcggcgcatgagagaagcccagaccaattacctacccaaaatggagaaagtagaagccaagcaggtcactgataatgaccatgctaatgccagagcgttttcgcatctggcttcaaaactggtgttaaatcattcagctacctgagaggggcccctataactctctacggctaacctgaatggagccaccatgttcgtgttcctggttctgttaccactggtgagcagccaatgtgtgaatctgacgaccagaacccagctgcctcctgcctacaccaatagctttacaaggggcgtgtattaccctgacaaggtctttcggagcagcgtcctgcacagcacccaggacctgttcttgcccttcttcagcaacgtgacctggttccacgccatacacgtgtccggcaccaatggcaccaagcgcttcgacaaccccgtgctgcctttcaacgacggcgtgtacttcgcctcaactgaaaagtccaatatcatccggggctggatcttcggcactacactggactctaagacccagagcctgctgatcgtgaacaatgccacaaacgtggtgattaaggtctgcgagttccagttctgtaacgaccctttcctgggagtgtactaccataagaacaacaagagctggatggaaagcgaattcagagtgtacagctctgctaataactgtaccttcgagtacgtgagccagcctttcctgatggacctggaaggcaagcagggcaacttcaagaacctgcgggaattcgtgttcaagaacatcgacggctacttcaaaatttactccaagcacacacctatcaacttggtgagagacctgccccagggctttagcgccctggaacctctggtggatctgcctatcggcatcaacatcaccagattccagaccctgctggcccttcacagatcctacctgactccaggcgacagctcctctggctggaccgccggcgccgctgcttactacgtcggatacctgcaacccagaaccttcctgctgaagtataacgagaacggcacaatcacagatgcggtggactgcgccctggaccccctgagcgagacaaagtgcaccctgaagtctttcaccgttgagaaaggcatctaccagacaagcaactttcgggtgcagcccaccgagagcatcgtgaggtttccaaacatcaccaacctgtgccctttcggagaagtgttcaacgccaccagattcgccagcgtgtacgcctggaacagaaagcggatcagcaactgtgtggctgactactctgtgctgtataactccgcgagtttcagcacctttaagtgctacggcgtgtcccccaccaagctgaacgacctctgcttcaccaacgtgtatgccgatagcttcgtaatccggggggacgaggtgcgacagatcgcccctggacagaccggaaacattgccgactacaactacaagctgcctgatgatttcactggatgcgtgatcgcctggaactccaacaacctggacagtaaagtgggcggaaattacaattacctgtacagactcttcagaaagagcaacctcaagccttttgaaagagatatcagcacagagatttaccaggccggcagcaaaccttgcaacggcgtggccggcttcaactgttactttcctctgcaaagctacggcttccagccaacatacggcgtgggctatcagccatacagagtggtggtgctgagcttcgagctgctccacgcacccgcgaccgtgtgcggccccaagaagtccaccaacctggtcaagaacaagtgcgtgaacttcaatttcaacggcctgactggcactggagtgctgaccgagagcaacaaaaagttcctgccattccaacagtttggtagagacatcgccgatacaacagacgccgtgagggatccccagaccttagaaatcctggatatcaccccttgttcatttggcggcgtgtctgtgatcacgcccggcaccaacaccagcaaccaggtggccgtgctgtaccaaggcgtcaattgcaccgaggtgcctgtggccatccacgccgaccagctgacccctacatggagagtgtacagcaccggcagcaacgtgttccagaccagagccggctgcctgatcggtgctgaatacgtgaataattcttacgagtgcgacatccctatcggcgccggcatctgcgccagctaccagacccagacaaattcccccggctctgcctcttctgttgcctcacagtccatcatcgcctacacaatgtctctgggcgctgaaaattccgtcgcttacagcaataacagcatcgccatccctacaaacttcactatcagcgttacaaccgagattctgcctgttagcatgaccaaaaccagcgtggattgcaccatgtacatctgcggcgacagcaccgagtgtagcaatctgctgctgcaatacggctccttttgtacccagctgaaccgggcgctgaccggaatcgctgtggaacaggacaagaacacccaggaggtgttcgcccaagtgaaacagatctataagacccctcctatcaaggacttcggcggatttaatttcagccagatcctgcctgacccttccaagcccagcaagcggtctcctatcgaggacctgctctttaacaaggtcaccctggctgatgccggcttcatcaagcagtacggcgattgcctgggcgatatcgccgctagagatctgatctgtgcccaaaaattcaacggcctgacagtgctgccccctctgctgactgacgagatgattgcccagtacactagcgccctgctggctggaacgatcacctctggctggaccttcggcgccggccctgccctgcaaatccccttccccatgcagatggcctacagattcaacggaatcggagtgacacaaaacgtgctgtacgagaaccagaagctcatcgctaaccagttcaacagcgccatcgggaagatccaggactcactcagcagcacacctagcgccctgggtaagctgcaagacgtagtgaaccagaacgcccaggccctgaacaccctggtgaagcagctgagttctaactttggagccatctcttccgtgctgaacgacatcctgagcagactggacccacctgaggccgaagtgcagatcgatcggctgatcacaggcagactccagtccctgcaaacctacgtgacacagcagctgatccgggccgccgagatcagagcctctgccaacctcgcagctacaaaaatgagcgagtgtgtgctgggccagagcaagcgggtggacttctgcggcaaaggctatcacctgatgagctttccccagagcgcccctcacggcgtcgtattcctgcacgtcacctacgtgcccgctcaagagaagaacttcaccacagcccctgccatctgccacgatggcaaggcccacttccccagagagggagtgtttgtcagcaacggcacgcactggttcgtgacccaaagaaacttctacgagcctcagatcatcacaaccgataataccttcgtgagcggaaattgcgacgtggtcatcggcatcgtgaacaacacagtgtacgaccccctgcaacctgaactggatagcttcaaggaggagctggacaagtacttcaaaaaccacaccagccctgatgtggacctgggcgacatcagcggcattaacgccagcgtggtgaatatccagaaggaaatcgacagactgaacgaagtggctaagaatctgaacgagtctctgattgacctgcaagagctgggcaaatacgagcagtacatcaaatggccttggtacatctggctgggcttcatcgccgggctgatcgccatcgtgatggtgaccatcatgctgtgctgcatgaccagctgctgtagctgcctgaagggctgctgtagctgcggcagctgctgtaaattcgacgaagatgactctgaacctgtgctgaagggcgtcaagctgcattacaccattttgtttttaatatttcaaaaaaaaaaaaaaaaaaaaaaaaaaaaaaaaaaaaaaaaaaaaaaaaaaaaaaaaaaaaaaaaaaaaaaaaaaaaaaaaaaaaaaaaaaaaaaaa

**4. *In vitro* confirmation of elevated SARS-CoV-2 spike protein expression in the designed TA mRNA vaccine**

**Fig.**
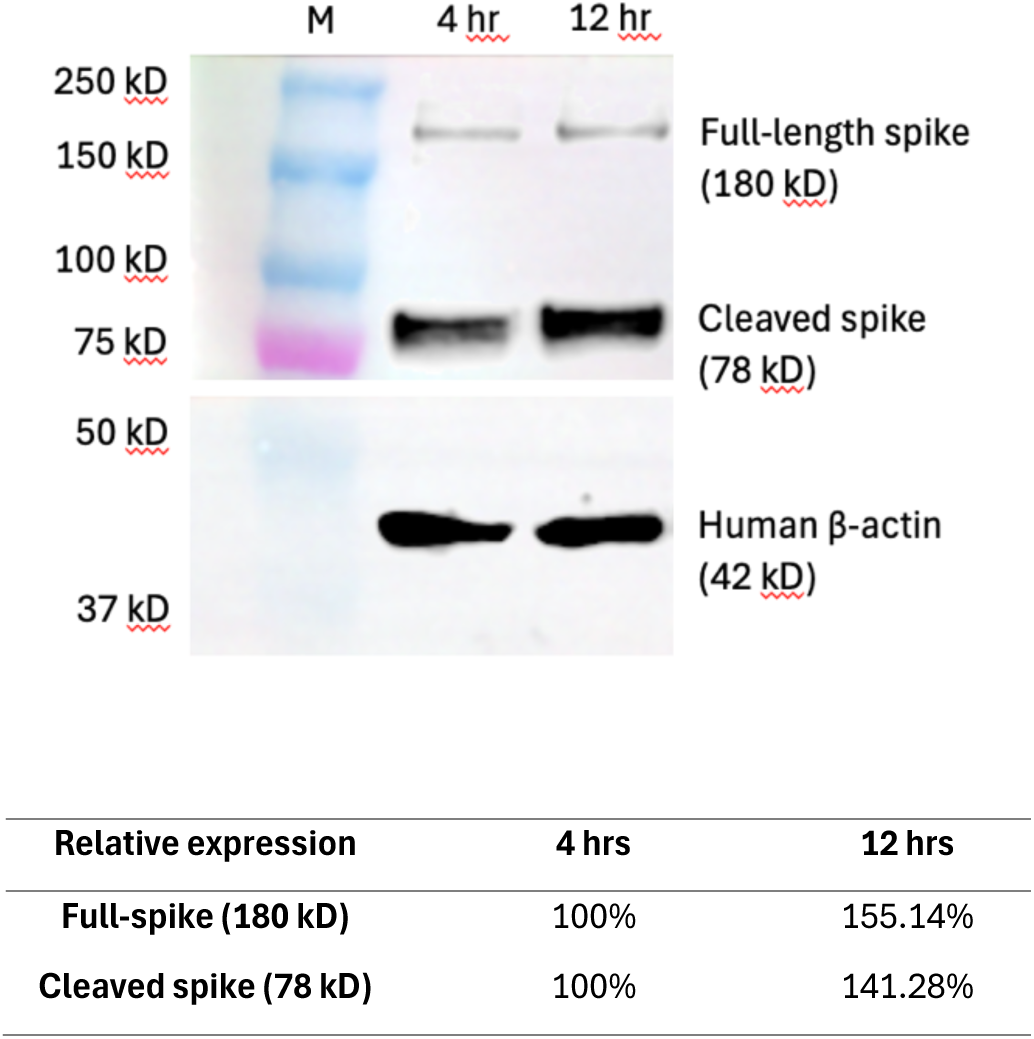
Western blot analysis of SARS-CoV-2 spike protein expression in A549 cells co-transfected with mRNAs encoding VEEV replicase and SARS-CoV-2 spike protein at 4 hours and 12 hours post-transfection. Spike protein expression increased over time, with higher levels of both the full-length (180 kDa, 155.14%) and cleaved (78 kDa, 141.28%) forms observed at 12 hours. Beta-actin (42 kDa) was used as a loading control. Lane 1: Marker; Lane 2: Cell lysate at 4 hours; Lane 3: Cell lysate at 12 hours.

**5. Reduced expression of IL-6 in the cells co-transfected with VEEV replicase and SARS-CoV-2 mRNA group compared to the control**

**Fig.**
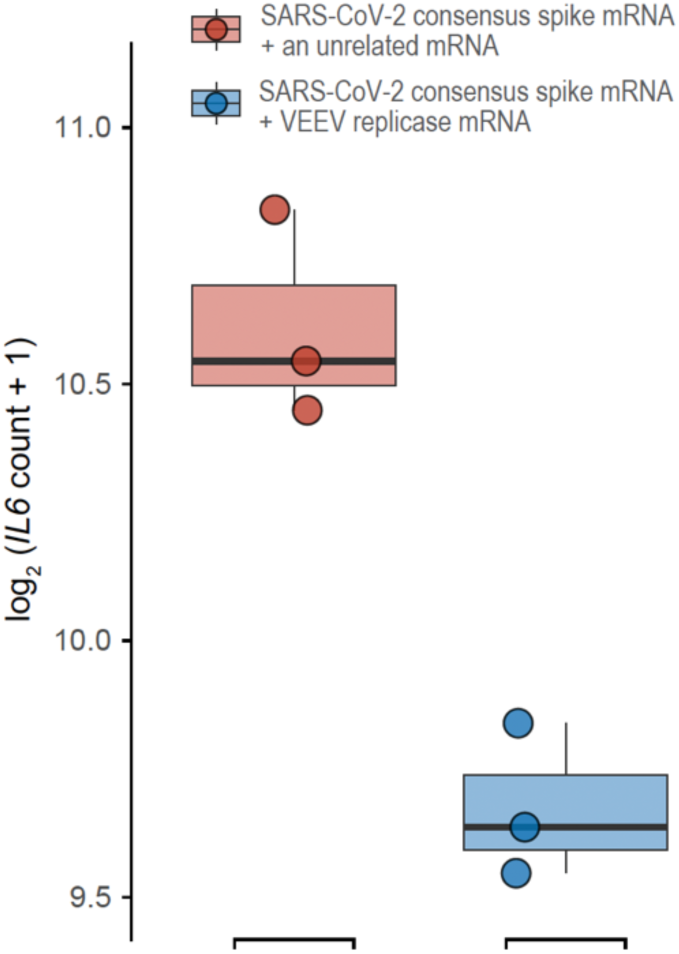
IL-6 expression in A549 cells 12 hours post mRNA transfection. Cells transfected with VEEV replicase mRNA and SARS-CoV-2 mRNA (treatment group, *n*=3) demonstrated a reduction in IL-6 expression compared to cells transfected with a SARS-CoV-2 mRNA and an unrelated mRNA (control group, *n*=3).

## References

1. Markov PV, et al. The evolution of SARS-CoV-2. Nature Reviews Microbiology 21, 361–379 (2023).

2. Beesley LJ, et al. SARS-CoV-2 variant transition dynamics are associated with vaccination rates, number of co-circulating variants, and convalescent immunity. EBioMedicine 91, 104534 (2023).

3. https://www.ecdc.europa.eu/en/covid-19/variants-concern. (2024).

4. https://stacks.cdc.gov/view/cdc/116721. (2022).

5. Khandia R, et al. Emergence of SARS-CoV-2 Omicron (B.1.1.529) variant, salient features, high global health concerns and strategies to counter it amid ongoing COVID-19 pandemic. Environ Res 209, 112816 (2022).

6. Aleem A, Akbar Samad AB, Vaqar S. Emerging Variants of SARS-CoV-2 and Novel Therapeutics Against Coronavirus (COVID-19). In: StatPearls). StatPearls Publishing, Copyright © 2023, StatPearls Publishing LLC. (2023).

7. Giovanetti M, et al. Evolution patterns of SARS-CoV-2: Snapshot on its genome variants. Biochemical and Biophysical Research Communications 538, 88–91 (2021).

8. Rochman ND, Wolf YI, Faure G, Mutz P, Zhang F, Koonin EV. Ongoing global and regional adaptive evolution of SARS-CoV-2. Proceedings of the National Academy of Sciences 118, e2104241118 (2021).

9. Carabelli AM, et al. SARS-CoV-2 variant biology: immune escape, transmission and fitness. Nature Reviews Microbiology 21, 162–177 (2023).

10. Yap C, et al. Comprehensive literature review on COVID-19 vaccines and role of SARS-CoV-2 variants in the pandemic. Therapeutic Advances in Vaccines and Immunotherapy 9, 25151355211059791 (2021).

11. Barouch DH. Covid-19 Vaccines — Immunity, Variants, Boosters. New England Journal of Medicine 387, 1011–1020 (2022).

12. Moore JP, Offit PA. SARS-CoV-2 Vaccines and the Growing Threat of Viral Variants. JAMA 325, 821–822 (2021).

13. Grant R, et al. When to update COVID-19 vaccine composition. Nature Medicine 29, 776–780 (2023).

14. Andeweg SP, et al. Elevated risk of infection with SARS-CoV-2 Beta, Gamma, and Delta variants compared with Alpha variant in vaccinated individuals. Science Translational Medicine 15, eabn4338 (2022).

15. Paul P, et al. Effectiveness of the pre-Omicron COVID-19 vaccines against Omicron in reducing infection, hospitalization, severity, and mortality compared to Delta and other variants: A systematic review. Hum Vaccin Immunother 19, 2167410 (2023).

16. Sarwar MU, et al. Real-world effectiveness of the inactivated COVID-19 vaccines against variant of concerns: meta-analysis. Journal of Infection and Public Health, (2023).

17. Chae C, et al. Comparing the effectiveness of bivalent and monovalent COVID-19 vaccines against COVID-19 infection during the winter season of 2022-2023: A real-world retrospective observational matched cohort study in the Republic of Korea. Int J Infect Dis 135, 95–100 (2023).

18. Gozzi N, et al. Estimating the impact of COVID-19 vaccine inequities: a modeling study. Nature Communications 14, 3272 (2023).

19. Chen J, Chen J, Xu Q. Current Developments and Challenges of mRNA Vaccines. Annual Review of Biomedical Engineering 24, 85–109 (2022).

20. Rosa SS, Prazeres DMF, Azevedo AM, Marques MPC. mRNA vaccines manufacturing: Challenges and bottlenecks. Vaccine 39, 2190–2200 (2021).

21. Echaide M, Chocarro de Erauso L, Bocanegra A, Blanco E, Kochan G, Escors D. mRNA Vaccines against SARS-CoV-2: Advantages and Caveats. Int J Mol Sci 24, (2023).

22. Vogel AB, et al. Self-Amplifying RNA Vaccines Give Equivalent Protection against Influenza to mRNA Vaccines but at Much Lower Doses. Mol Ther 26, 446–455 (2018).

23. Beissert T, et al. A Trans-amplifying RNA Vaccine Strategy for Induction of Potent Protective Immunity. Mol Ther 28, 119–128 (2020).

24. Bloom K, van den Berg F, Arbuthnot P. Self-amplifying RNA vaccines for infectious diseases. Gene Therapy 28, 117–129 (2021).

25. Fuller DH, Berglund P. Amplifying RNA Vaccine Development. New England Journal of Medicine 382, 2469–2471 (2020).

26. McCafferty S, et al. A dual-antigen self-amplifying RNA SARS-CoV-2 vaccine induces potent humoral and cellular immune responses and protects against SARS-CoV-2 variants through T cell-mediated immunity. Mol Ther 30, 2968–2983 (2022).

27. Palladino G, et al. Self-amplifying mRNA SARS-CoV-2 vaccines raise cross-reactive immune response to variants and prevent infection in animal models. Molecular Therapy - Methods & Clinical Development 25, 225–235 (2022).

28. Voigt EA, et al. A self-amplifying RNA vaccine against COVID-19 with long-term room-temperature stability. npj Vaccines 7, 136 (2022).

29. Schmidt C, et al. A taRNA vaccine candidate induces a specific immune response that protects mice against Chikungunya virus infections. Mol Ther Nucleic Acids 28, 743–754 (2022).

30. Dai L, Gao GF. Viral targets for vaccines against COVID-19. Nature Reviews Immunology 21, 73–82 (2021).

31. Lee K-M, Lin S-J, Wu C-J, Kuo R-L. Race with virus evolution: The development and application of mRNA vaccines against SARS-CoV-2. Biomedical Journal 46, 70–80 (2023).

32. Cohen KW, et al. Trivalent mosaic or consensus HIV immunogens prime humoral and broader cellular immune responses in adults. J Clin Invest 133, (2023).

33. Mezhenskaya D, Isakova-Sivak I, Rudenko L. M2e-based universal influenza vaccines: a historical overview and new approaches to development. Journal of Biomedical Science 26, 76 (2019).

34. Wang R, et al. Vaccination With a Single Consensus Envelope Protein Ectodomain Sequence Administered in a Heterologous Regimen Induces Tetravalent Immune Responses and Protection Against Dengue Viruses in Mice. Front Microbiol 10, 1113 (2019).

35. Zuo L, et al. Design and Characterization of a DNA Vaccine Based on Spike with Consensus Nucleotide Sequence against Infectious Bronchitis Virus. Vaccines (Basel) 9, (2021).

36. Muthumani K, et al. A synthetic consensus anti-spike protein DNA vaccine induces protective immunity against Middle East respiratory syndrome coronavirus in nonhuman primates. Sci Transl Med 7, 301ra132 (2015).

37. Hsieh CL, et al. Structure-based design of prefusion-stabilized SARS-CoV-2 spikes. Science 369, 1501–1505 (2020).

38. Cromer D, et al. Predicting COVID-19 booster immunogenicity against future SARS-CoV-2 variants and the benefits of vaccine updates. Nature Communications 15, 8395 (2024).

39. Cromer D, et al. Neutralising antibody titres as predictors of protection against SARS-CoV-2 variants and the impact of boosting: a meta-analysis. The Lancet Microbe 3, e52–e61 (2022).

40. Gilbert Peter B, Donis Ruben O, Koup Richard A, Fong Y, Plotkin Stanley A, Follmann D. A Covid-19 Milestone Attained — A Correlate of Protection for Vaccines. New England Journal of Medicine 387, 2203–2206 (2022).

41. Ahmed WS, Philip AM, Biswas KH. Decreased Interfacial Dynamics Caused by the N501Y Mutation in the SARS-CoV-2 S1 Spike:ACE2 Complex. Front Mol Biosci 9, 846996 (2022).

42. Liu Y, et al. The N501Y spike substitution enhances SARS-CoV-2 infection and transmission. Nature 602, 294–299 (2022).

43. Miotto M, et al. Inferring the stabilization effects of SARS-CoV-2 variants on the binding with ACE2 receptor. Communications Biology 5, 20221 (2022).

44. Calvaresi V, et al. Structural dynamics in the evolution of SARS-CoV-2 spike glycoprotein. Nat Commun 14, 1421 (2023).

45. Gellenoncourt S, et al. The Spike-Stabilizing D614G Mutation Interacts with S1/S2 Cleavage Site Mutations To Promote the Infectious Potential of SARS-CoV-2 Variants. J Virol 96, e0130122 (2022).

46. Gobeil SM, et al. D614G Mutation Alters SARS-CoV-2 Spike Conformation and Enhances Protease Cleavage at the S1/S2 Junction. Cell Rep 34, 108630 (2021).

47. Korber B, et al. Tracking Changes in SARS-CoV-2 Spike: Evidence that D614G Increases Infectivity of the COVID-19 Virus. Cell 182, 812–827.e819 (2020).

48. Zhou B, et al. SARS-CoV-2 spike D614G change enhances replication and transmission. Nature 592, 122–127 (2021).

49. Yurkovetskiy L, et al. S:D614G and S:H655Y are gateway mutations that act epistatically to promote SARS-CoV-2 variant fitness. bioRxiv, (2023).

50. Rutten L, et al. Impact of SARS-CoV-2 spike stability and RBD exposure on antigenicity and immunogenicity. Sci Rep 14, 5735 (2024).

51. Zhao Y, et al. Vaccination with S(pan), an antigen guided by SARS-CoV-2 S protein evolution, protects against challenge with viral variants in mice. Sci Transl Med 15, eabo3332 (2023).

52. Chen M-W, et al. A consensus–hemagglutinin-based DNA vaccine that protects mice against divergent H5N1 influenza viruses. Proceedings of the National Academy of Sciences 105, 13538–13543 (2008).

53. Olvera A, Noguera-Julian M, Kilpelainen A, Romero-Martín L, Prado JG, Brander C. SARS-CoV-2 Consensus-Sequence and Matching Overlapping Peptides Design for COVID19 Immune Studies and Vaccine Development. Vaccines (Basel) 8, (2020).

54. Sun H, Sur J-H, Sillman S, Steffen D, Vu HLX. Design and characterization of a consensus hemagglutinin vaccine immunogen against H3 influenza A viruses of swine. Veterinary Microbiology 239, 108451 (2019).

55. Xia X. Detailed Dissection and Critical Evaluation of the Pfizer/BioNTech and Moderna mRNA Vaccines. Vaccines (Basel) 9, (2021).

56. Schmidt C, et al. A Bivalent Trans-Amplifying RNA Vaccine Candidate Induces Potent Chikungunya and Ross River Virus Specific Immune Responses. Vaccines (Basel) 10, (2022).

57. Lello LS, et al. Cross-utilisation of template RNAs by alphavirus replicases. PLOS Pathogens 16, e1008825 (2020).

58. Strauss JH, Strauss EG. The alphaviruses: gene expression, replication, and evolution. Microbiol Rev 58, 491–562 (1994).

59. Orlandini von Niessen AG, et al. Improving mRNA-Based Therapeutic Gene Delivery by Expression-Augmenting 3’ UTRs Identified by Cellular Library Screening. Mol Ther 27, 824–836 (2019).

60. Bhalla N, Sun C, Metthew Lam LK, Gardner CL, Ryman KD, Klimstra WB. Host translation shutoff mediated by non-structural protein 2 is a critical factor in the antiviral state resistance of Venezuelan equine encephalitis virus. Virology 496, 147–165 (2016).

61. Simmons JD, et al. Venezuelan equine encephalitis virus disrupts STAT1 signaling by distinct mechanisms independent of host shutoff. J Virol 83, 10571–10581 (2009).

62. Pardi N, Hogan MJ, Porter FW, Weissman D. mRNA vaccines — a new era in vaccinology. Nature Reviews Drug Discovery 17, 261–279 (2018).

63. Wroblewska L, et al. Mammalian synthetic circuits with RNA binding proteins for RNA-only delivery. Nat Biotechnol 33, 839–841 (2015).

64. Holtkamp S, et al. Modification of antigen-encoding RNA increases stability, translational efficacy, and T-cell stimulatory capacity of dendritic cells. Blood 108, 4009–4017 (2006).

65. Halfmann PJ, et al. SARS-CoV-2 Omicron virus causes attenuated disease in mice and hamsters. Nature 603, 687–692 (2022).

66. Frische A, et al. Optimization and evaluation of a live virus SARS-CoV-2 neutralization assay. PLoS One 17, e0272298 (2022).

67. Crawford KHD, et al. Protocol and Reagents for Pseudotyping Lentiviral Particles with SARS-CoV-2 Spike Protein for Neutralization Assays. Viruses 12, (2020).

68. Cho H, et al. Bispecific antibodies targeting distinct regions of the spike protein potently neutralize SARS-CoV-2 variants of concern. Sci Transl Med 13, eabj5413 (2021).

69. Dacon C, et al. Broadly neutralizing antibodies target the coronavirus fusion peptide. Science 377, 728–735 (2022).

70. Yuan M, et al. Structural and functional ramifications of antigenic drift in recent SARS-CoV-2 variants. Science 373, 818–823 (2021).

71. Zhou P, et al. Broadly neutralizing anti-S2 antibodies protect against all three human betacoronaviruses that cause deadly disease. Immunity 56, 669–686.e667 (2023).

72. Ritz C, Baty F, Streibig JC, Gerhard D. Dose-Response Analysis Using R. PLOS ONE 10, e0146021 (2016).

73. Kumar S, et al. Selection of Ideal Reference Genes for Gene Expression Analysis in COVID-19 and Mucormycosis. Microbiol Spectr 10, e0165622 (2022).

74. Lu X, et al. US CDC Real-Time Reverse Transcription PCR Panel for Detection of Severe Acute Respiratory Syndrome Coronavirus 2. Emerg Infect Dis 26, 1654–1665 (2020).

75. Reed LJ, Muench H. A simple method of estimating fifty per cent endpoints. American journal of epidemiology 27, 493–497 (1938).

76. Andrews S. FastQC: A Quality Control Tool for High Throughput Sequence Data. https://www.bioinformatics.babraham.ac.uk/projects/fastqc/. (2010).

77. Chen S, Zhou Y, Chen Y, Gu J. fastp: an ultra-fast all-in-one FASTQ preprocessor. Bioinformatics 34, i884–i890 (2018).

78. Kim D, Paggi JM, Park C, Bennett C, Salzberg SL. Graph-based genome alignment and genotyping with HISAT2 and HISAT-genotype. Nature Biotechnology 37, 907–915 (2019).

79. Love MI, Huber W, Anders S. Moderated estimation of fold change and dispersion for RNA-seq data with DESeq2. Genome Biology 15, 550 (2014).

80. Liao Y, Smyth GK, Shi W. featureCounts: an efficient general purpose program for assigning sequence reads to genomic features. Bioinformatics 30, 923–930 (2014).

81. Yu G, Wang LG, Han Y, He QY. clusterProfiler: an R package for comparing biological themes among gene clusters. Omics 16, 284–287 (2012).

82. Kassambara A. ggpubr: ‘ggplot2’ Based Publication Ready Plots. R package version 0.6.0. (2023).

